# Surfactin facilitates the establishment of *Bacillus subtilis* in synthetic communities

**DOI:** 10.1101/2024.08.14.607878

**Authors:** Carlos N. Lozano-Andrade, Caja Dinesen, Mario Wibowo, Nil Arenos Bach, Viktor Hesselberg-Thomsen, Scott A. Jarmusch, Mikael Lenz Strube, Ákos T. Kovács

## Abstract

Soil bacteria are prolific producers of a myriad of biologically active secondary metabolites. These natural products play key roles in modern society, finding use as anti-cancer agents, as food additives, and as alternatives to chemical pesticides. As for their original role in interbacterial communication, secondary metabolites have been extensively studied under *in vitro* conditions, revealing a multitude of roles including antagonism, effects on motility, niche colonization, signaling, and cellular differentiation. Despite the growing body of knowledge on their mode of action, biosynthesis, and regulation, we still do not fully understand the role of secondary metabolites on the ecology of the producers and resident communities *in situ.* Here, we specifically examine the influence of *Bacillus subtilis*-produced cyclic lipopeptides (LPs) during the assembly of a bacterial synthetic community (SynCom), and simultaneously, explore the impact of LPs on *B. subtilis* establishment success in a SynCom propagated in an artificial soil microcosm. We found that surfactin production facilitates *B. subtilis* establishment success within multiple SynComs. Surprisingly, while neither a wild type nor a LP non-producer mutant had major impact on the SynCom composition over time, the *B. subtilis* and the SynCom metabolomes are both altered during co-cultivation. Overall, our work demonstrates the importance of surfactin production in microbial communities, suggesting a broad spectrum of action of this natural product.

## Introduction

Microbes produce a plethora of small molecules with diverse activities, which are extensively exploited in modern society. A number of these natural products, often denoted as secondary or specialized metabolites (SMs), have been pivotal in contemporary medicine and biotechnological industries [1, 2]. They serve as frontline therapy against infectious diseases, therapeutics for cancer [1], food additives [3], or crop protection agents [4]. Besides the long-standing tradition of industrial exploitation, SMs are considered chemical mediators that modulate interactions within and between microbial species or even cross-kingdoms. For instance, defensive molecules might help producers defend their resources or niche from microbial competitors [5]. Furthermore, some SMs function as signal molecules for coordinated growth (i.e. for quorum-sensing) [6, 7] and cell-differentiation [8, 9].

Despite growing knowledge about their mode of action, biosynthesis, and regulation, the natural functions exerted by these metabolites are still not well understood in natural systems in an ecological context [10–13]. Here, we aimed to fill this gap by exploring the role of cyclic lipopeptides (LPs) produced by a *Bacillus subtilis* isolate [14–16] during bacterial synthetic community (SynCom) assembly, and simultaneously, dissecting the impact of LPs on *B. subtilis* establishment success in a SynCom.

The *B. subtilis* species complex encompasses a diverse group of soil-dwelling strains with prolific potential for SM production, including lipopeptides, polyketides, ribosomally synthesized and post-transcriptionally modified peptides, and signaling molecules [14, 17–20]. Among these SMs, cyclic lipopeptides (LPs) are the most widely studied class. They are synthesized by non-ribosomal peptide synthase (NRPS), acting as a molecular assembly line that catalyzes the incorporation of amino acids into a growing peptide [21]. LPs in the *B. subtilis* species group are structurally clustered into three families: surfactins, iturins and fengycins, according to their peptide core sequence. They are composed of seven (surfactins and iturins) or ten (fengycins) α-amino acids linked to β-amino (iturins) or β-hydroxyl (surfactin and fengycins) fatty acids [22, 23]. LPs are an example of multifunctional SMs; they can act as antimicrobials by antagonizing other microorganisms [14, 24–26] but also play pivotal roles in processes including motility, surface colonization, signaling, and cellular differentiation [27–30].

Most evidence supporting the multifaceted role of LPs has been gathered under *in vitro* conditions with pure cultures, which may not reflect the complexity of soil environments and community-level interactions modulating the function of these SMs. Moreover, the high diversity of soil microbiomes and the difficulty in tracking the *in situ* production of LPs and other classes of specialized metabolites have made these questions difficult to address experimentally [11, 19]. As a consequence, the use of less complex systems mimicking natural conditions has emerged as a useful approach to disentangle the inherent complexity of interactions mediated by SMs *in situ* [31, 32], and namely the use of synthetic bacterial communities (SynCom) has been a promising strategy for exploring fundamental patterns in controlled, but ecological relevant, conditions [33, 34]. For instance, Cairns *et. al* used a 62-strain SynCom to demonstrate how low antibiotic concentration impacts community composition and horizontal transfer of resistance genes, while Niu et. al built a seven-member community mimicking the core microbiome of maize, which was able to protect the host from a plant-pathogenic fungus [35, 36]. Simultaneously to the establishment of the concept of SynComs, the development of soil-like matrices and artificial soil has provided a useful option for studying chemical ecology in highly controlled gnotobiotic systems compatible with analytical chemistry and microbiological methods [37–39]. Thus, coupling the use of artificial soil system and simplified SynCom is a fast-growing approach to examine microbial interactions, while maintaining some degree of ecological complexity.

In this study, we aimed to reveal the role of LPs in two ecological processes: the ability of the LP-producer *B. subtilis* to thrive in a SynCom and the influence of such LP-producing bacterium on SynCom assembly. To address this, an artificial soil-mimicking system [40, 41] was used to assess the impact of non-ribosomal peptides (s*fp*), and specifically surfactin (*srfAC*) or pliplastatin (*ppsC*) on the ability of *B. subtilis* to establish within a four-member SynCom. We demonstrated that surfactin production facilitates *B. subtilis* establishment success within SynCom in a soil-mimicking environment. Surprisingly, the wild-type and non-producer strains had a comparable influence on the SynCom composition over time. Moreover, we revealed that the *B. subtilis* and SynCom metabolome are both altered. Intriguingly, the importance of surfactin for the establishment of *B. subtilis* has been demonstrated in diverse SynCom systems with variable composition. Altogether, our work expands the knowledge about the role of surfactin production in microbial communities, suggesting a broad spectrum of action of this natural product.

## Results

### Description of the artificial soil system inoculated with SynCom

To assess the role of *B. subtilis* SMs on bacterial community assembly under soil-mimicking conditions, we have previously customized a hydrogel matrix that allows us to grow several bacterial strains axenically and quantify the specific *B. subtilis* lipopeptides (*i.e.* surfactin and plipastatin) [41]. In addition, we assembled a four-membered bacterial synthetic community (SynCom) isolated from the same sample site as *B. subtilis* P5_B1 [42]. In addition to the shared origin with *B. subtilis*, we selected these four isolates since the co-existence in our hydrogel beads system and their morphological distinctness allowed us to quantify them by plate count readily. Although the relative abundance of each of the four strains fluctuated throughout the experiments, all four members were still detectable for up to six days of sampling (Fig 1A). At the end of the experiment, we observed a clear strain co-existence pattern in the SynCom as previously observed: *Stenothrophomonas indicatrix* and *Chryseobacterium* sp. were the most dominant strains, *Rhodococcus globerulus* was kept at low density whereas *Pedobacter* sp. was outcompeted at 14 days (Fig S1). Using this established experimental system, we explored the role of LPs in the successful establishment of *B. subtilis*, as well as in SynCom assembly and functionality. A schematic diagram illustrating the core experimental design, and the scientific question is presented in Figure 1.

**Figure 1.**
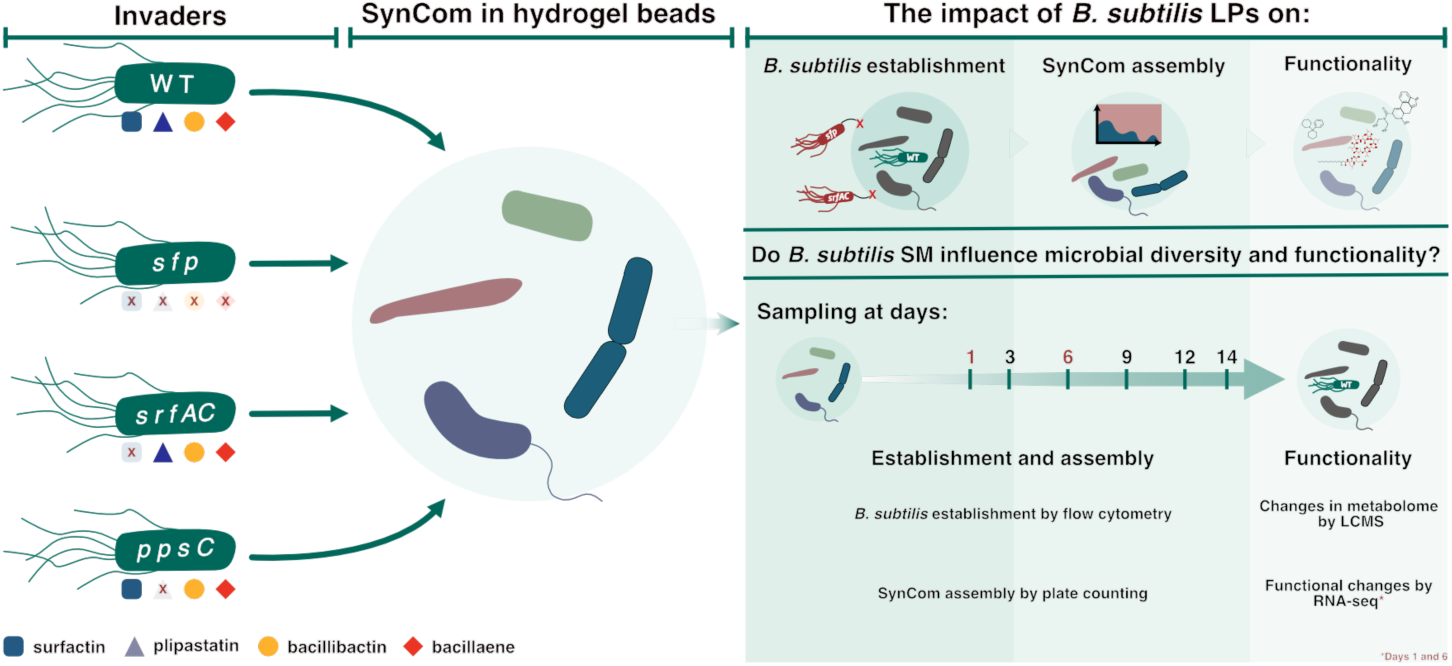
A schematic diagram illustrating the experimental design, and the research question of the main experiments conducted. *B. subtilis* P5_B1 and its NRP-deficient mutants were inoculated into a 4-member SynCom propagated in the hydrogel bead microcosms.

### Surfactin production facilitates *B. subtilis* P5_B1 establishment in a four-member SynCom

To evaluate the contribution of specific lipopeptides to P5_B1 establishment in the SynCom, we co-cultivated either the WT strain or the SM production-impaired mutants (*sfp*, *srfAC* and *ppsC*)in the presence of the SynCom using the hydrogel matrix that mimics soil characteristics [40, 41]. Initially, we confirmed that P5_B1 and its mutant derivatives grew and produced the expected lipopeptides when cultivated axenically in the soil-like system. All *B. subtilis* strains colonized the hydrogel system at comparable rates (ANOVA at day 14, *P=0.87*), demonstrating a similar population dynamic pattern: a one-log increase within a day followed by a plateau of nearly 1x10^7^ CFU/g of the hydrogel after three days of cultivation, which was maintained up to the final sampling time on day 14 (Figure 2C).

When introduced to the SynCom, the WT and *ppsC* mutant (that produce surfactin but not plipastatin), successfully colonized the beads and maintained their population at approximately 1x10^7^ CFU/g throughout the experiment, comparable to the titers obtained in axenic cultivation. In contrast, the population size of the *B. subtilis* genotypic variants impaired in non-ribosomal peptides (*sfp*) or solely in surfactin (*srfAC*) production sharply declined during the first six days. By the end of the experiment, the cell titers decreased to around three log-fold below the initial population levels (ANOVA, P < 0.01) (Figure 2B). Following up on these observations, we investigated whether the WT strain could rescue the *srfAC* mutant by co-inoculating a mixture of both strains into the SynCom. In this co-culture, the WT strain remained more competitive than the *srfAC* mutant. However, the presence of the WT strain, and presumable its surfactin production capability, evidently rescued the *srfAC* mutant, as its decline was less pronounced compared to when introduced alone into the SynCom (Fig 1D).

**Figure 2.**
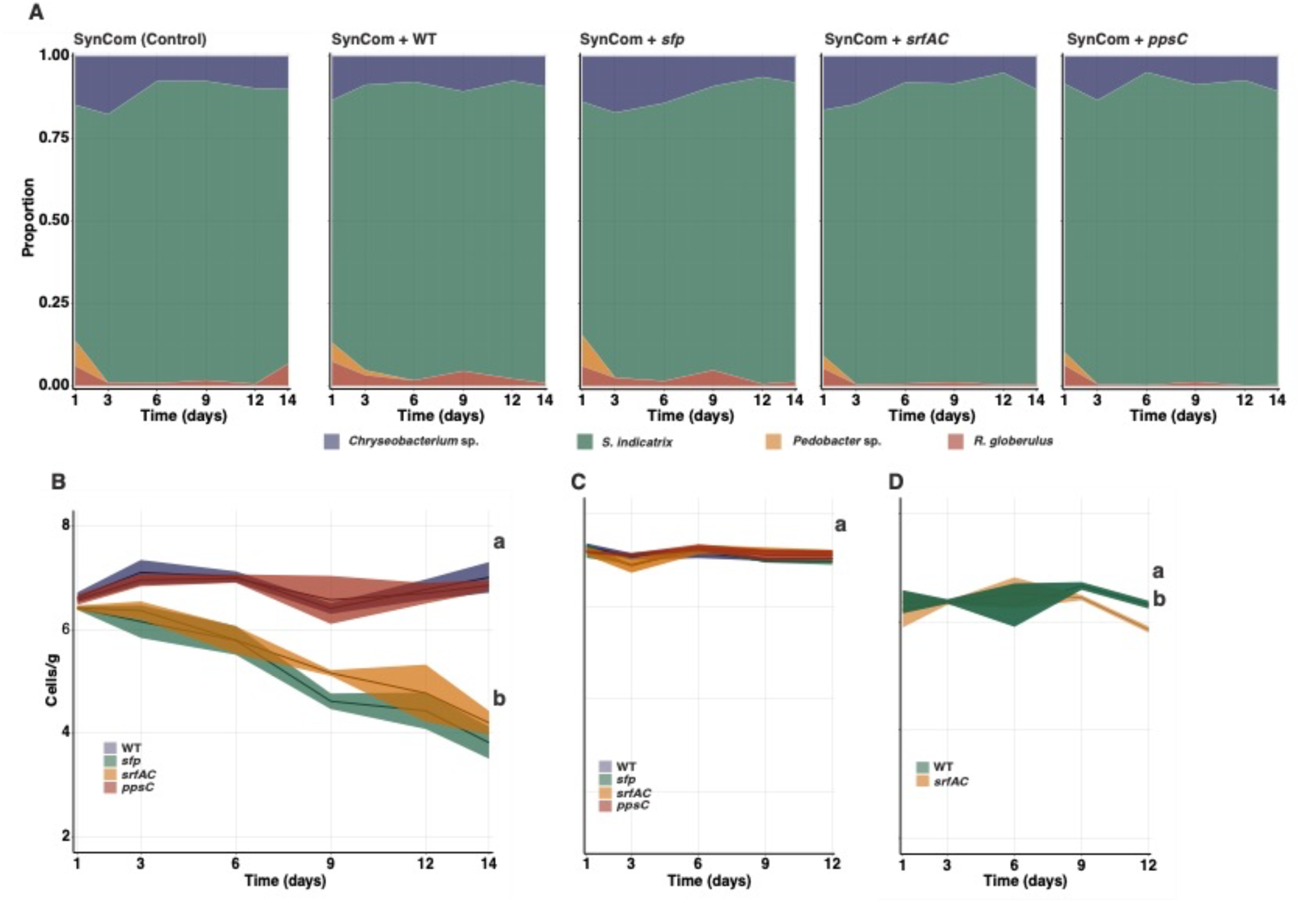
Surfactin production facilitates *B. subtilis* establishment in a SynCom but does not alter its composition in the soil-like environment over time. A set of *gfp*-labeled *B. subtilis* strains (Surfactin producers: WT and *ppsC*, non-producers: *sfp* and *srfAC*) were introduced into the SynCom, and their populations were followed over time to determine the role of LPs in SynCom assembly and their contribution to *B. subtilis* establishment in the simplified system. (A). SynCom assembly after the different *B. subtilis* variants were introduced. Members abundances are represented as the proportion occupied by each member relative to the total biomass in the system. (B) *B. subtilis gfp*-labeled population dynamics after introduction in the SynCom. Surfactin producers (WT and *ppsC*) population size remains stable over time while non-producers (*sfp* and *srfAC*) sharply decline by the end of the experiments suggesting that surfactin production is crucial for *B. subtilis* establishment in the SynCom. C) *B. subtilis* WT and its derivates mutant growth dynamics when propagated individually in the hydrogel microcosms. D) Complementation assay. A mixed population (1:1) of the WT and the *srfAC* was propagated in the hydrogel beads in the presence of the SynCom. The presence of the WT strain (surfactin producer) rescued the surfactin-deficient mutant. The letters represent significant differences among groups at day 14 (one-way ANOVA and Tukey Honest tests). The experiment was conducted independently twice with n=3 in both cases. The data were pooled and analyzed together.

Subsequently, we investigated the potential contribution of individual SynCom members to the decline of the surfactin-deficient strains using a *pair-wise* competition assay in planktonic cultures. Here, varying ratios of each SynCom member and *B. subtilis* were assessed and the reduction of the growth (i.e. area under the curve) relative to the monoculture was measured. *B. subtilis* populations experienced a significant reduction when co-cultured with *S. indicatrix* D763 and *Chryseobacterium* sp. D764 at the highest ratio (1, 0.1, 0.01 of the tested strain relative to the *B. subtilis* cultures), irrespective of *B. subtilis* capability to produce surfactin. However, in co-cultures where the SynCom members were diluted (more than 0.01 relative to *B. subtilis*), *B. subtilis* strains lacking surfactin production were outcompeted by *S. indicatrix* D763 and *Chryseobacterium* sp. D764. Overall, *B. subtilis* WT showed greater competitiveness against these SynCom members, maintaining higher growth at higher dilution ratios compared to the *sfp* and *srfAC* mutants. In contrast, the less competitive strains in the bead systems, *R. globerulus* D757 and *Pedobacter* sp. D749, only impacted *B. subtilis* growth at the highest co-culture ratio, with strains lacking surfactin production exhibiting comparable growth to WT (Figure 3).

**Figure 3.**
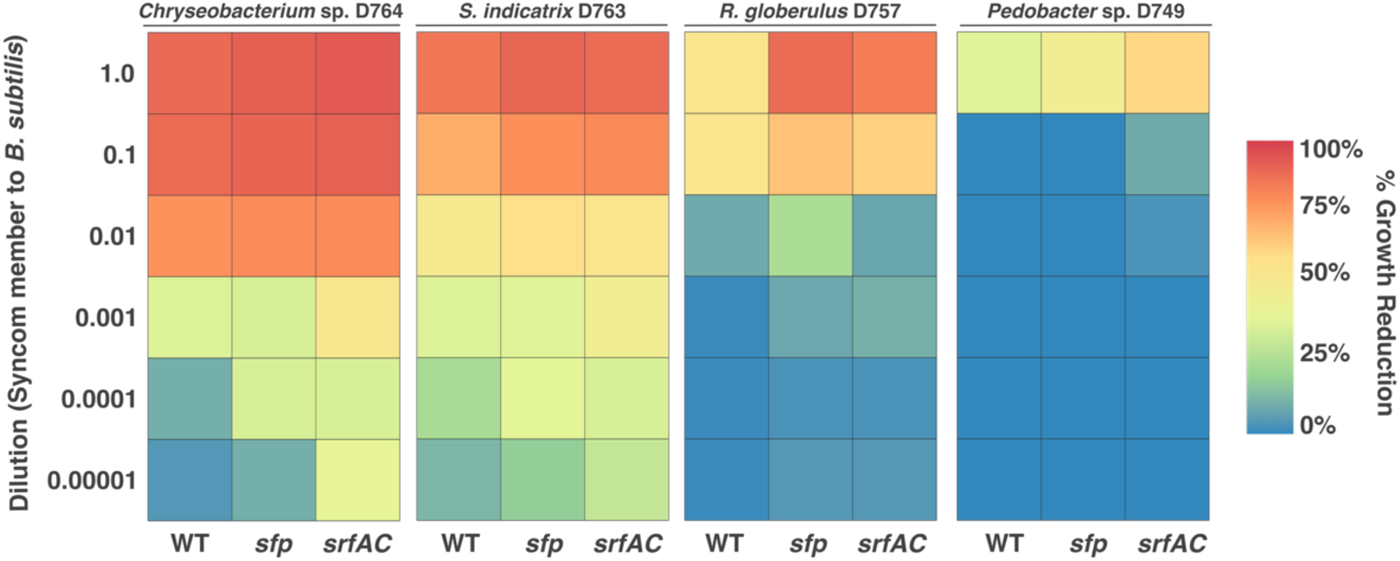
Impact of individual SynCom members on *B. subtilis* growth. A set of *B. subtilis* strains were co-cultured with each SynCom member at different ratios. The GFP signal was used as a proxy for the respective *B. subtilis* strains’ growth in co-culture and the area under the curve of growth was used as culture productivity parameter. The impact of being co-culture with each SynCom member was estimated as %*Growth reduction = [(Growth BsubMonoculture − Growth Bsub Co − culture)/Growth Bsbu Monoculture)] × 100*.

### *B. subtilis* secondary metabolites do not have a major impact on SynCom assembly

Motivated by our observation that SM production, specifically surfactin, plays a crucial role in *B. subtilis* establishment success, we investigated if these SMs impact the SynCom composition over time. To do this, we evaluated the abundance of SynCom members (CFU) using NMDS and PERMANOVA (Fig 4). Regardless of the *B. subtilis* strain introduced, the SynCom followed similar assembly dynamics as we described above: *S. indicatrix* and *Chryseobacterium* sp. dominated the community while *R. globerulus* and *Pedobacter* sp. were less abundant (Figure 2A and Fig S2). Estimation of the growth rates and the carrying capacity of each SynCom member in 0.1× TSB revealed that *S. indicatrix*, the most dominant strain, grew significantly faster and reached the highest productivity whereas *Pedobacter* sp. grew at the slowest rate (Fig S3). This could explain the observed SynCom composition on the hydrogel system, which was dominated by the fastest growers and more productive strains.

**Fig 4.**
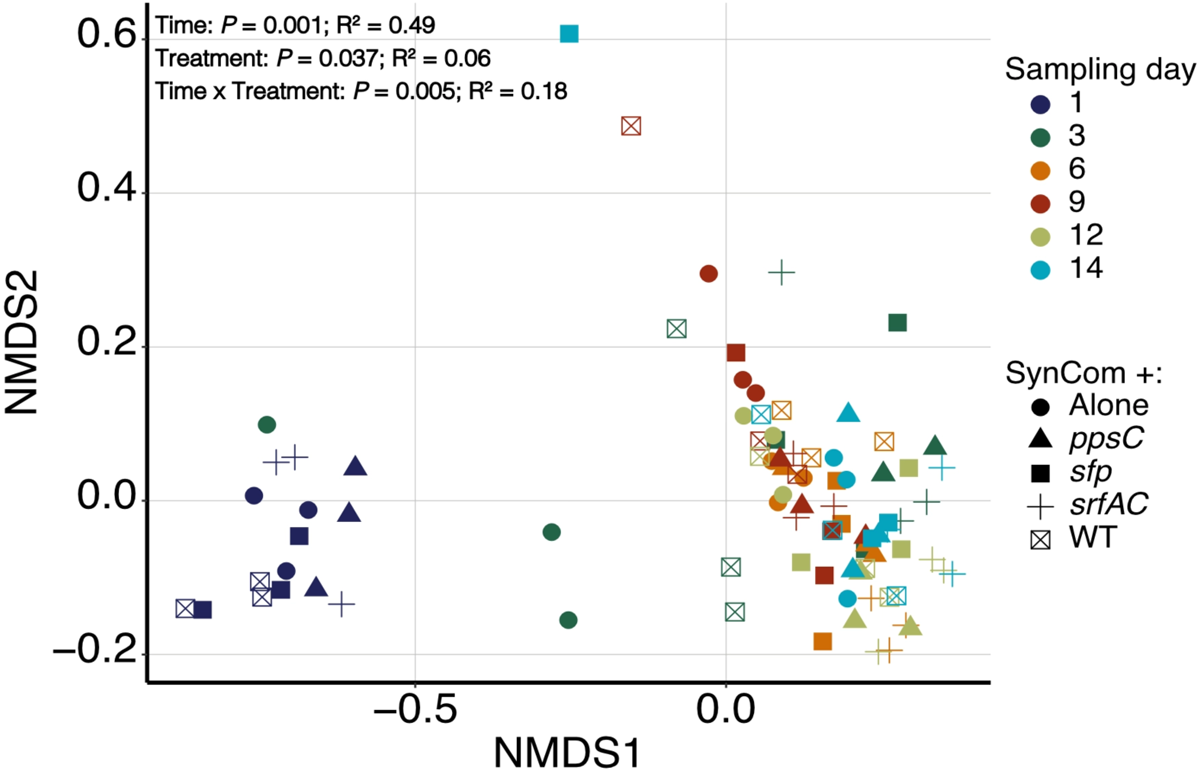
Changes in SynCom composition upon *B. subtilis* inoculation. Bray-Curtis distance NMDS ordination plot performed on the CFU data of the SynCom after *B. subtilis* introduction for comparing the effects of Sampling time (colors) and the variant of *B. subtilis* (shapes) on SynCom composition. The statistical differences were assessed by a PERMANOVA. The *P* values and R^2^ are reported as an inset within the figure.

A fixed-effect PERMANOVA using sampling time, *B. subtilis* variants and their interaction confirmed that the main driver of SynCom composition was the sampling time (PERMANOVA, R^2^ = 0.49, *P* = 0.001), with a minor effect of *B. subtilis* strain introduced (PERMANOVA, R^2^ = 0.06, *P* = 0.037) and the interaction (PERMANOVA, R^2^ = 0.18, *P* = 0.005). Overall, the results suggested that introducing either the WT or its SM-impaired mutants did not have a major impact on the SynCom assembly, with the differences mainly explained by the sampling time (Fig 4).

Next, we investigated whether the antagonistic activity between the SynCom members and *B. subtilis* could explain our observations. Using an *in vitro* inhibition test, we found that the less competitive strains, *Pedobacter* sp. D749 and *R. globerulus* D757, were both susceptible to *B. subtilis*. Specifically, the activity of *Pedobacter* sp. D749 was linked to NRP production, particularly surfactin, while *R. globerulus* was inhibited by all the variants. This suggests that other classes of SMs beyond NRP, produced by *B. subtilis*, might be accounted for inhibition of these two species. Nevertheless, the SynCom-abundant strains, *S. indicatrix* D763 and *Chryseobacterium* sp. D764, displayed no growth reduction by *B. subtilis* and its SMs, as evidenced by the absence of an inhibition zone (Fig S4).

### *B. subtilis* and SynCom metabolome are both altered during the establishment experiments

To explore the role of *B. subtilis* secondary metabolites in shaping the SynCom metabolome and how surfactin production was modulated in co-cultivation, we profiled both the SynCom and *B. subtilis* metabolome at day 14 of the experiment using liquid chromatography-mass spectrometry (LC-MS). A targeted approach revealed that the production of surfactin was significantly increased when the WT variant was grown in the presence of the SynCom compared with the WT production in axenic cultures (t-test, *P* = 0.0317) (Figure 5A). This finding was further validated *in vitro* by supplementing P5_B1 cultures with cell-free supernatants from each of the SynCom members or all strains together. Interestingly, the spent media from both the monocultures and the SynCom induced the production of surfactin, with the highest increase observed when P5_B1 was supplemented with *R. globerulus* supernatant (Figure 5B).

**Figure 5.**
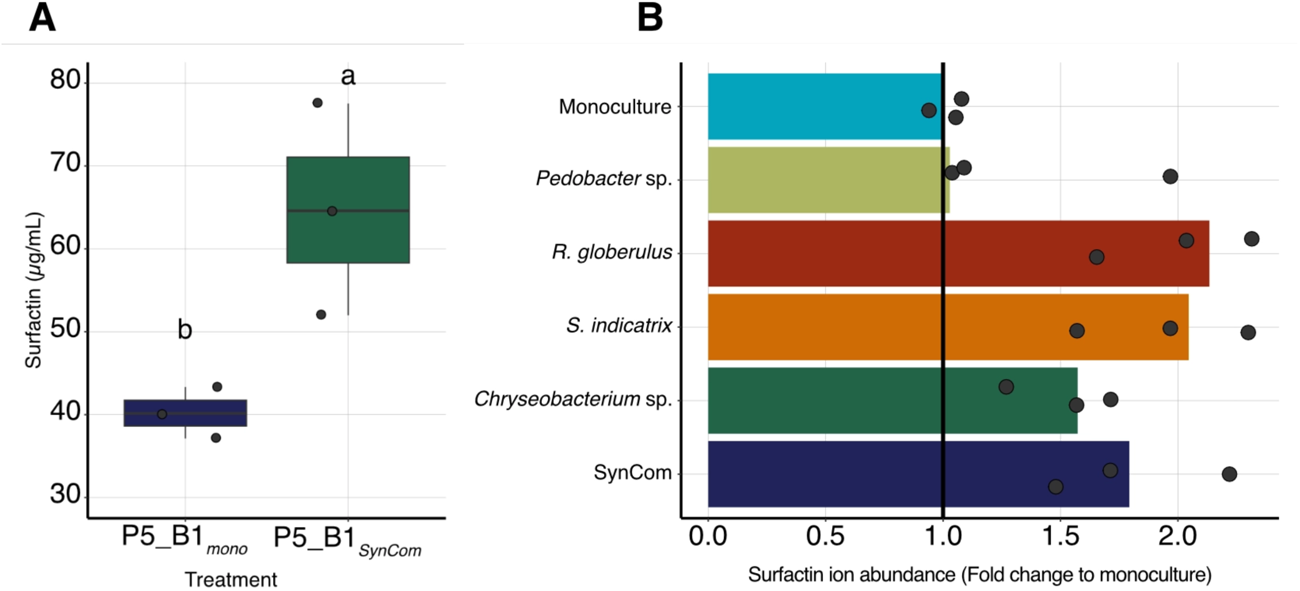
The full SynCom and individual SynCom members induce surfactin production both in the bead system and in liquid culture. (A) Surfactin production in the bead system by P5_B1 increased when co-cultivated with the full SynCom compared to pure *B. subtilis* monoculture. The concentration was estimated from the last day of sampling in the *B. subtilis* – SynCom co-cultivation experiment. (B) Changes in surfactin production when P5_B1 was supplemented with cell-free supernatants obtained from individual SynCom members and the full SynCom in liquid cultures. In both experiments surfactin production was quantified by UHPLC and three replicates (n = 3) were performed per treatment.

Moreover, untargeted analysis of the SynCom upon co-inoculation of surfactin-producing *B. subtilis* revealed extensive metabolic changes suggesting that surfactin production alters SynCom chemical landscape (Figure 6A). When we determined the number of features detected in each community over time, three clear patterns emerged from the data: stable and slowly increasing chemical output (SynCom+*sfp*), steady decline to day 6 and then increase at day 9 (SynCom+WT and SynCom), and rapid decline at day 3 followed by steady increase to day 9. The SynCom challenged with the WT strain has the highest overall chemical diversity at day 1 and 9. Without the products of NRP biosynthesis (in the *sfp* mutant), the SynCom system seems more compatible and does not undergo a metabolic change like seen in the other systems (Fig S5).

Although most of the molecular features (*m/z*) detected in our system remained unidentified, the molecular network clearly shows the presence of the *B. subtilis* lipopeptides, plipastatin, and surfactin, and their analogs. Moreover, the presence of ornithine lipids (OLs) was observed in the dataset [43]. These metabolites are derived from Gram negative bacterial cell outer membrane as surrogates of phospholipids under phosphate-limited conditions [44] (Fig S10). The lipid abundances (m/z between 597 and 671) increased in the SynCom alone, indicating this conversion of phospholipids to ornithine lipids occurs in the absence of *B. subtilis* (Figure 6). Ecologically, OLs have been linked to stress response [43]. Interestingly, when surfactin producers (WT or *ppsC* mutant) were introduced into the system, the presence of OLs was strongly reduced. Contrary, in the presence of the *sfp* and *srfAC* mutants, the OLs remain at levels comparable to the SynCom alone.

**Figure 6.**
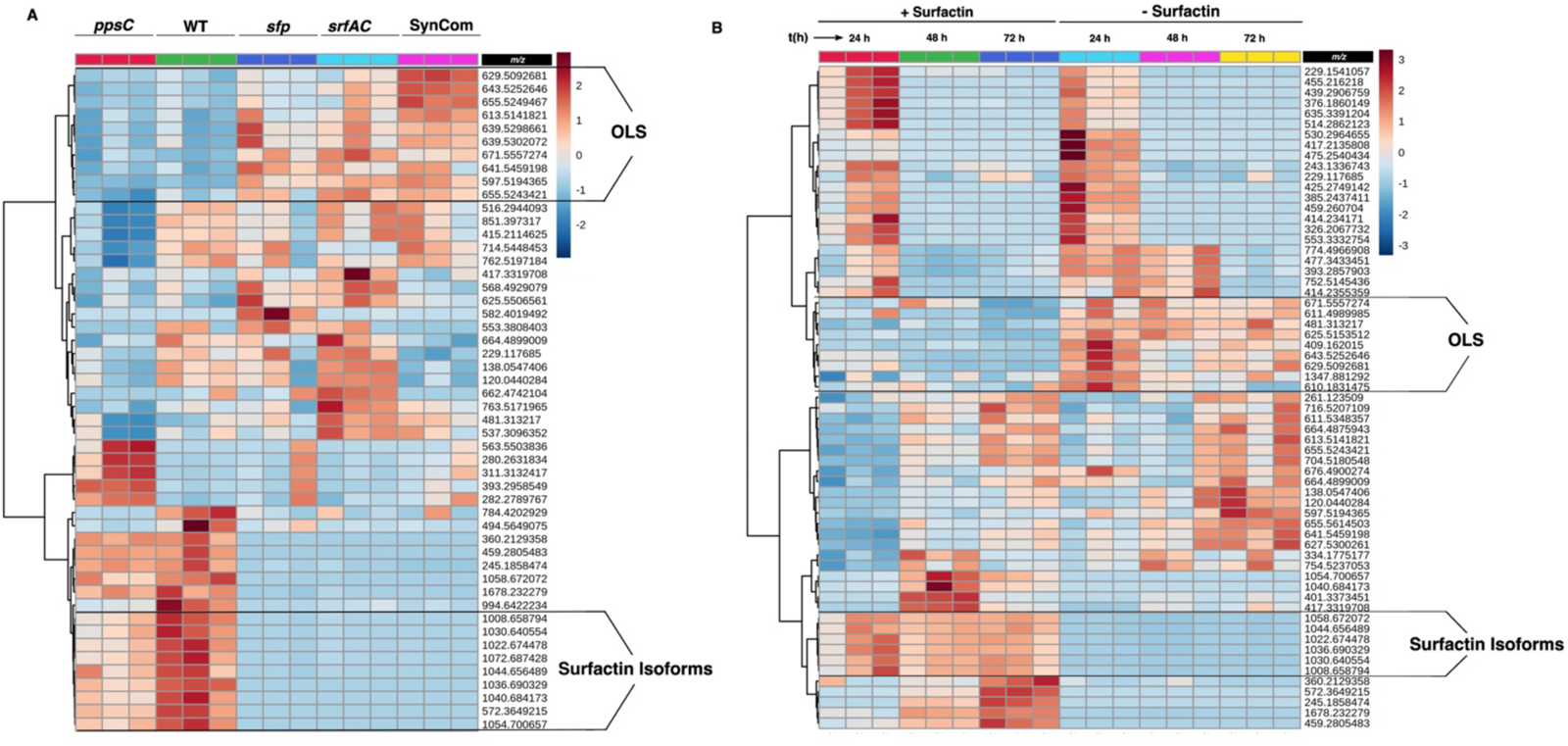
Untargeted metabolic analysis revealed extensive metabolic changes in the co-cultivation experiments. (A) A heatmap visualization based on the significantly increased or decreased chemical features (m/z) in the co-cultivation experiments. A set of chemical features (m/z), related to ornithine lipids (OLS) were strongly modulated in presence of surfactin producer (WT and *pps*C) (Upper brackets). Similarly, m/z associated with surfactin isoforms were detected in the system (Bottom brackets). (B) Changes in the SynCom chemical features when supplemented with pure surfactin. The visualization shows the variation over the time (24, 48 and 72 h) of the main chemical features produced by the SynCom. Similarly, the same group, OLS-lipids, were produced less in presence of pure surfactin (upper brackets). The heatmaps were made on the feature abundances retrieved from the ESI-MS chromatogram.

We corroborated this observation by conducting an experiment with the SynCom in the presence of pure surfactin. Here, the same group of compounds (*m/z* features) was altered in the surfactin-supplemented SynCom culture as in the presence of surfactin-producing *B. subtilis* co-cultures, while these were abundant in the control samples (*i.e*., without *B. subtilis*) (Figure 6B).

### The less competitive strains of the SynCom were the most transcriptionally affected species by *B. subtilis* SMs

To dissect the mechanism of how surfactin facilitates *B. subtilis* establishment within the SynCom, a meta-transcriptomic approach was conducted comparing the transcriptional profile of the SynCom challenged with the WT and the *sfp* mutant. In total, 430 genes and 490 genes were differentially expressed (DEG) in the SynCom after 1 and 5 days, respectively, inoculated with the WT compared with the sample seeded with the *sfp* mutant. In both sampling days, the less competitive strains, *Pedobacter* sp. D749 and *R. globerulus* D757 had the highest number of differentially expressed genes (DGEs) in the system, accounting for around the 83% of DEGs at day 1 and 95% of those at the last sampling point (Fig S11). Subsequently, we explored the distribution of clusters of orthologous groups (COG categories) among the DEGs genes to discover which processes within the SynCom are potentially affected by the introduction of either the WT or *sfp* mutant. Here, many DEGs were not annotated or classified as COG S, an unknown function. However, cell wall/membrane/envelope biogenesis (COG M) and amino acid transport and metabolism (COG E) were the most abundant functional categories among the genes downregulated in the SynCom with WT strain added relative to the SynCom in the presence of *sfp* mutant (Fig S12).

Additionally, we explored the functions and enrichment pathways of DEGs for the less competitive strains (*Pedobacter* sp. D749 and *R. globerulus* D757). The gene ontology (GO) enrichment analysis revealed that both strains responded transcriptionally different in the presence of the WT strains. While the enriched biological processes in *R. globerulus* D757 were related to defense mechanisms or response to other organisms, upregulated processes in *Pedobacter* sp. were linked to amino acid transport, specifically histidine (Fig S13)

### Surfactin-facilitated establishment of *B. subtilis* is remarkably conserved across diverse SynComs

To survey if surfactin is important for establishment of *B. subtilis* P5_B1 within diverse microbial communities, we assessed the abundance of WT and surfactin-deficient mutant in five previously published and characterized SynComs [36, 45–50]. These SynComs varied in composition, reflecting different functionalities and ecological niches. Overall, the co-culture experiments revealed that the ability of *B. subtilis* to establish within the SynComs depended on surfactin production, SynCom composition (number of members) and the inoculation ratio. In most SynComs, except for the Kolter Lab’s SynCom which was broadly invaded, both the WT and the *srfAC* mutant displayed reduced growth at a high inoculation ratio of SynCom (10:1, 1:1, 1:10). However, the WT, which produces surfactin, generally reached higher population densities compared to the surfactin-deficient mutant across most SynComs. When *B. subtilis* was inoculated at high ratios relative to the SynComs, the growth dynamics resembled those observed in axenic cultures of both the WT and *srfAC* mutant (Figure 7).

**Figure 7.**
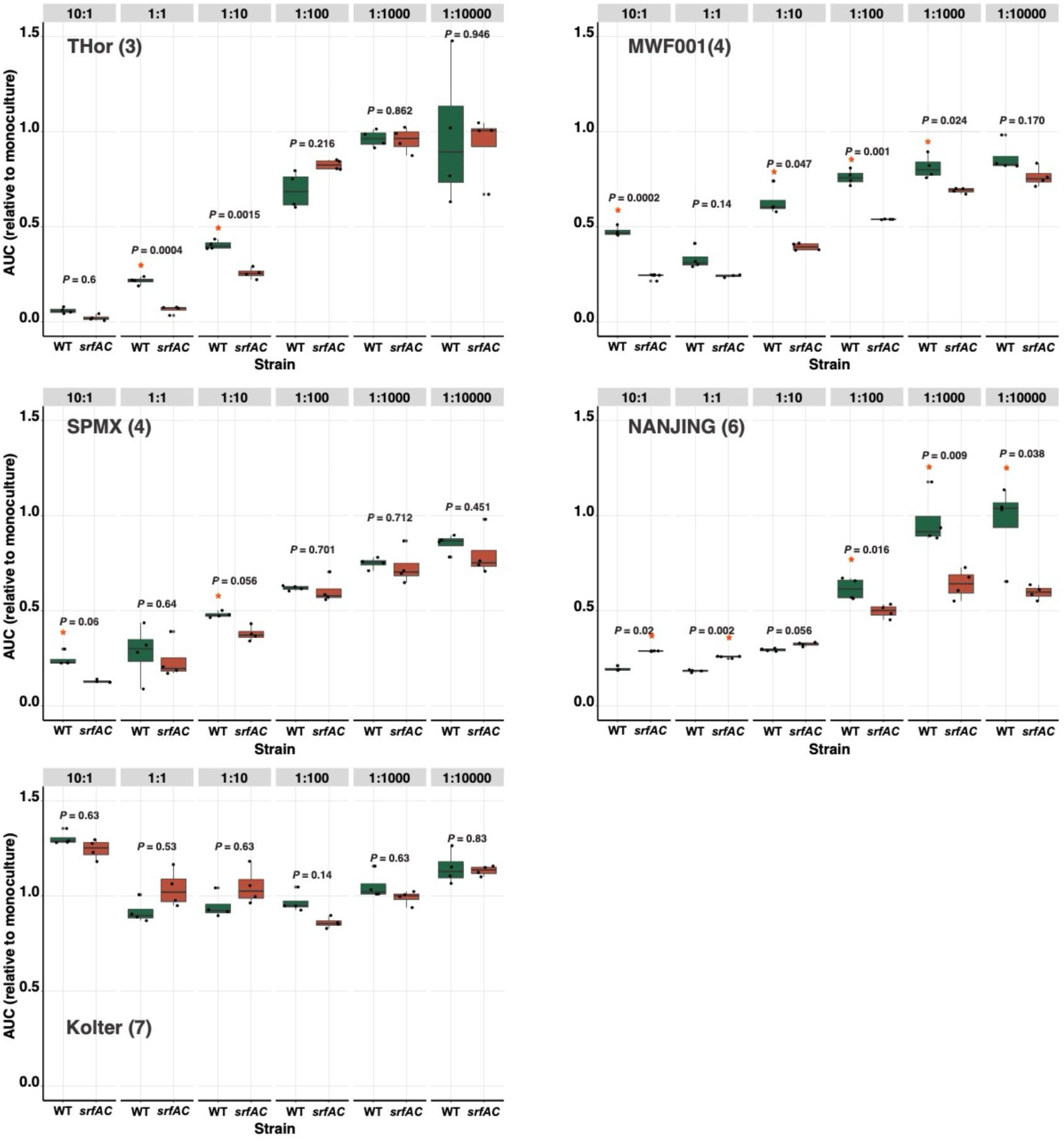
*B. subtilis* P5_B1 establishment in different publicly avalaible SynComs. The WT and its derivate mutant impaired in surfactin production (*srfAC*) were co-cultured with different SynCom at decreasing ratios (see Methods and SI for details). The *gfp* signal was used as proxy for *Bacillus* establishment in the co-cultures during 24h incubation period. The area under the curve (AUC) ratio (*B. subtilis* growth in co-culture in each SynCom dilution to *B. subtilis* growth in monoculture) was used as *B. subtilis* establishment parameter. “*” indicates that the value of the AUC ratio was significant and its position denotated which strain had a higher value. “ns” indicates not significance (*P*<0.05, Student’s t-test adjusted for multiple testing by the Benjamini-Hochberg method). The name inset the plot indicates the lab or the name of the SynCom, in parenthesis the number of members composing the community.

## Discussion

Traditionally, secondary metabolites have been mostly studied because of their antimicrobial or anticancer properties. However, several of these natural products exert a multifaceted function, influencing the physiology of the producing microorganism and modulating interactions with other organisms [10, 12]. Determining the role of these compounds in natural habitats, *e. g.* in soil, is challenging, given the chemical and biological complexity, and the limitations for SMs quantification *in situ*. This study was initiated to dissect the role of *B. subtilis*-produced LPs during SynCom assembly and their impact on the producers in a soil-like environment. We demonstrated that surfactin production facilitates *B. subtilis* establishment success across multiple SynComs, modulates the SynCom chemodiversity but does not alter the community composition.

We experimentally demonstrated the contribution of surfactin in *B. subtilis* success when inoculated in the presence of a SynCom using a reductionist approach: four-member bacterial SynCom propagated in microcosms based on an artificial hydrogel matrix [41, 42]. One of the biggest methodological challenges in studying SM-driven microbial interactions is to mimic the environmental conditions. Consequently, the need for developing model systems of intermediate complexity for elucidating the ecological role of these molecules and shedding light on microbiome assembly-related questions has been widely stated [31, 33, 51–53]. This is because classic axenic *in vitro* assays do not resemble crucial aspects of microbial niches, whereas natural samples are far too complex to dissect the underlying processes at the molecular level. Our SynCom is not intended to represent the natural sample site, i.e. Dyrehaven soil community, where all strains used in this study were isolated, but rather, it represents a reproducible, trackable, and easy-to-set bacterial assemblage useful for testing the role of SMs in SynCom assembly, and together with the soil-mimicking matrix, might help to overcome the bottlenecks imposed by soil complexity in terms of microbial diversity and SMs quantification. The described system aligns conceptually with recent approaches that used transparent microcosm mimicking the complexity of natural environments while testing hypotheses with statistical power in a controlled set up [40, 54–56].

Throughout the present work, we revealed that surfactin production is crucial for *B. subtilis* establishment success in a set of diverse SynComs. Surfactin is by far one of the most-studied LPs produced by *B. subtilis*. The relevance of this multifunctional SM has been demonstrated in biofilm formation [28, 57], swarming and sliding motility[58–62], root and phyllosphere colonization [63, 64], and triggering induced systemic resistance (ISR) in plants [65, 66] . Although it is not frequently highlighted as a primary function of surfactin, its contribution to producers’ fitness has been showed in different environmental conditions. For instance, Luo *et al* demonstrated that a *B. subtilis* strain impaired in surfactin production did not colonize rice sheaths inoculated with *R. solani.* At the same time, WT increased its population size over time [67]. Similarly, Zeriouh and colleagues showed that *srfAB* mutant (of *Bacillus amyloliquefaciens* UMAF6614) presents reduced persistence in the melon phylloplane[68]. In soil, similar observations were made where surfactin-impaired mutants of *B. subtilis* were unable to colonize *Arabidopsis thaliana* roots [30, 64]. In all these examples, the underlying mechanism is the link between surfactin production and bacterial differentiation (surface spreading and colonization). Even though further experiments are required to provide a mechanistic explanation of why and how surfactin facilities *B. subtilis* establishment in the SynCom, *B. subtilis* P5_B1 wild type and its surfactin mutant derivative have been demonstrated to produce biofilm *in vitro* and on plant roots under laboratory settings[28]. Alternatively, surfactin production could help *B. subtilis* to cope with a potential oxygen depletion induced by the SynCom growth. Such novel function of surfactin has been recently demonstrated where surfactin mediated *B. subtilis* survival via membrane depolarization and increased oxygen diffusion under low oxygen concentration[69].

Surprisingly, the WT and the SM-mutant strains had hardly any influence on the composition and dynamics of the SynCom, but surfactin production altered the chemical diversity of the SynCom, besides the sensitivity of minor SynCom members to *B. subtilis* SMs. Several studies are highlighted that isolates of the *B. subtilis* species complex are not strong competitors of indigenous soil microbiota and as consequence they did not shift the composition rhizosphere bacterial community in a considerable degree[70, 71] or mainly influenced specific group of the rhizospheres’ microbial community[72, 73] . However, application of *B. subtilis* and its close relative species in the rhizosphere improve plant health and resiliency, and SM production contributes to these properties.

Beyond the impact of the examined lipopeptides on *B. subtilis* growth dynamics and SynCom composition, we evaluated how *B. subtilis* – SynCom co-cultivation altered SM induction in *B. subtilis* and the SynCom-derived metabolites. Interestingly, surfactin production was enhanced in the presence of the SynCom or specific SynCom members compared to *B. subtilis* monocultures. Similarly, the SynCom-secreted metabolome was strongly modulated by the surfactin-producing *B. subtilis*. These observations align with the well-supported idea that competitive interactions may play a pivotal role in the production of antibiotics [74–78]. Certain *Bacillus* strains display phenotypic changes in the presence of SMs that could be considered as a defensive response upon sensing bacterial competitors. For instance, increased biofilm formation [57, 79, 80], enhanced motility [81], induction of sporulation[82], and secondary metabolites production [83, 84] are all potential defense response of these bacteria. Nevertheless, the underlying mechanisms behind specific regulation of *B. subtilis* SM production in response to their neighbor’s activity are often unknown. The so-called “competition sensing” hypothesis formulated the underlying ecological concept, suggesting that microbes have evolved the ability to sense a hazard signal coupled with a stress response that allows a “counterpunch” by upregulating the production of antibiotics and toxins [13, 85].

In sum, soil bacteria are well known for their potential to synthesize a plethora of SMs with a wide diversity of activities. Our understanding of the ecological roles of these metabolites under natural conditions has just begun to be unlocked. Our observations, gathered in an intermediate ecological complex experimental system revealed the role of surfactin in the ecology of the producers and how this SM impacts the metabolism of its interacting partners. Thus, we hypothesize that the production of multimodal secondary metabolites by *B. subtilis* is a refined strategy that contributes to fitness and persistence in natural habitats where competition could be thorough.

## Materials and methods

### Bacterial strains and culture media

All the strains used in this study are listed in Table S1. *B. subtilis* strains were routinely grown in lysogeny broth (LB) medium supplemented with the appropriated antibiotic (LB-Lennox, Carl Roth; 10 g·l^-1^ tryptone, 5 g·l^-1^ yeast extract, and 5 g·l^-1^ NaCl) at 37 °C with shaking at 220 rpm. The strains composing the different synthetic communities were grown in 0.5 × Trypticase Soy Broth (TSB; Sigma-Aldrich, St. Louis, Missouri, USA) for 24h at 28 °C with shaking at 220 rpm.

### *Bacillus* subtilis establishment in the Dyrehaven SynCom propagated in soil-like matrix

The impact of introducing *B. subtilis* P5_B1 and its secondary-metabolite-deficient mutants into the SynCom was investigated using an artificial soil-mimicking microcosm [41]. Spherical beads were created by dripping a polymer solution, comprising 9.6 g/l of Phytagel^TM^ and 2.4 g/l sodium alginate in distilled water, into a 2% CaCl2 cross-linker solution [40]. After soaking in 0.1× TSB as a nutrient solution for two hours, the beads were sieved to remove any residual medium. Twenty milliliters of beads were then transferred to 50 mL Falcon tubes. Cultures of *B. subtilis* P5_B1 and the four SynCom members were grown as described above. The members of the SynCom were mixed at different OD, since fast-growing strains (i.e. *S. indicatrix* and *Chryseobacterium* sp.) had to be included at low density to ensure SynCom stability. Specifically, *Pedobacter* sp. and *R. globerulus* were adjusted to OD 2.0, while *S. indicatrix* and *Chryseobacterium* sp. were adjusted to OD 0.1 before mixing. Suspensions of *B. subtilis* P5_B1 and its mutants were standardized to OD 2.0. Next, bacterial inocula were prepared by mixing equal volumes of these adjusted cultures (four members plus each *B. subtilis* strain, respectively), and 2 mL of this suspension was then inoculated into freshly prepared beads. The bead microcosms were statically incubated at room temperature. Concurrently, microcosms inoculated with each strain as a monoculture were set as controls. At days 1, 3, 6, 9, 12, and 14, one gram of beads was transferred into a 15 mL Falcon tube, diluted in 0.9% NaCl, and vortexed for 10 minutes at maximum speed to disrupt the beads. The suspensions were then used for cell number estimation via colony-forming unit (CFU) and flow cytometry. For colony counting, 100 µL of the sample was serially diluted, spread onto 0.1× TSA, and CFU were estimated after 3 days. For the quantification of *B. subtilis* using flow cytometry, the samples were first passed through a Miracloth (Millipore) to remove any trace of beads and diluted 100-fold in 0.9% NaCl. Subsequently, 1 mL of each sample was transferred to an Eppendorf tube and assayed on a flow cytometer (MACSQuant® VYB, Miltenyi Biotec). GFP-labeled *B. subtilis* was detected using the blue laser (488 nm) and filter B1 (525/50 nm). Controls with non-inoculated beads and 0.1× TSB were employed to identify background autofluorescence. Single events were gated into the GFP vs. SSC-A plot, where GFP-positive cells were identified for each sample.

### WT:*srfAC* complementation assay

Overnight cultures of the strains of interest (OD600 = 2.0; WT::mKate and *srfAC*::gfp) were premixed at a 1:1 ratio. The inoculum was prepared by mixing equal volumes of the premixed *Bacillus* suspension with each member of the SynCom. Subsequently, 2 mL of this mixture were inoculated into freshly prepared beads. Propagation of the microcosms and *B. subtilis* quantification were performed as described above.

### Detection of secondary metabolites from artificial soil microcosms

To extract secondary metabolites from the bead samples, 1g of bead was transferred into a 15 mL with 4 mL of isopropyl alcohol:ethyl acetate (1:3 v/v), containing 1% formic acid. The tubes were sonicated for 60 minutes and centrifuged at 13400 rpm for 3 minutes. Then, the extracts were evaporated under N_2_ overnight, re-suspended in 300 µL of methanol and centrifuged at 13400 rpm. The supernatants were transferred to HPLC vial and subjected to ultrahigh-performance liquid chromatography-high resolution mass spectrometry (UHPLC-HRMS) analysis. The running conditions and the subsequent data analysis were performed as previously described [41].

### Metatranscriptomic analysis

For the RNA sequencing, the SynCom was propagated in the artificial soil matrix and challenged with either *B. subtilis* P5_B1 or the mutant impaired in NRP synthesis (*sfp* mutant). A SynCom without *B. subtilis* inoculation served as the control group. On days 1 and 6, 4 g of beads from each treatment were snap-frozen in liquid nitrogen and stored at -80°C. The RNA extraction was performed using the RNeasy PowerSoil Total RNA Kit (QIAGEN) following the manufacturer’s instructions. After extraction, the samples were treated with the TURBODNA-free kit (ThermoFisher) to degrade the remaining DNA. The library preparation and sequencing were carried out by Novogene Europe on Illumina NovaSeq 6000 S4 flow cell with PE150.

The reads were demultiplexed by the sequencing facility. Subsequently, reads were trimmed using Trimmomatic v.0.39[86]. Quality assessment was performed using FASTQC, and reads were sorted with SortMeRNA v.4.2.0[87] to select only the non-rRNA reads for the downstream analysis. Reads were then mapped onto the genomes of the strains (D764, D763, D757, D749, and B. subtilis P5_B1) using Bowtie v.2-2.3.2 [88]. Differential gene expression analysis was conducted using the R package DESeq2[89] using the shrunken log2 fold change values for analysis [90] The p values of each gene were corrected using Benjamini and Hochberg’s approach for controlling the false discovery rate (FDR). A gene was considered as differentially expressed when absolute log2 fold change was greater than 2 and FDR was less than 0.05. For functional analysis, the protein-coding sequences were mapped with KEGG Ontology, Gene Ontology (GO) terms, and **Clusters of Orthologous Genes** (COGs) using eggNOG-mapper[91]. Then, the eggNOG-mapper annotated dataset was used for gene set enrichment for pathway analysis in GAGE [92]. Transcriptomic analysis was performed from three independent replicates for each sample.

### Inhibition assay

The *in vitro* antagonistic effect of *B. subtilis* P5_B1 and its secondary metabolite-deficient mutants was assessed using double-layer agar plate inhibition assays against each SynCom member (target bacterium). All strains were cultured for 24 hours in 0.1× TSB medium as described previously. The cultures underwent two washes with 0.9% NaCl followed by centrifugation at 10,000 rpm for 2 minutes, and OD_600_ was adjusted to 0.1. For the first layer, 10 mL of 0.1× TSA (1.5% agar) were poured into petri dishes and allowed to dry for 30 minutes. Then, 100 µL of each target bacterium was added to 10 mL of 0.1× TSB containing 0.9% agar preheated to 45°C. This mixture was evenly spread on top of the 0.1× TSA and dried for an additional 30 minutes. Subsequently, 5 µL of each *B. subtilis* suspension was spotted on each plate. The plates were then incubated at room temperature, followed by examination of the inhibition zones on the lawn formed in the top layer.

Similarly, we investigated the impact of exometabolites produced by SynCom members on the growth properties of *B. subtilis strains*. Spent media from SynCom cultures were collected after 48 hours of growth in 0.1× TSB at 25°C and 250 rpm, filtered through 0.22 µm filters, and stored at 4°C. Growth curves were generated in 96-well microtiter plates. Each well contained 180 µL of 0.1× TSB supplemented with 5% spent media from each SynCom strain and 20 µL of either *B. subtilis* WT or its mutants. Control wells contained only 0.1× TSB medium without spent media supplementation. Cultivation was carried out in a Synergy XHT multi-mode reader at 25°C with linear continuous shaking (3 mm), monitoring optical density at 600 nm every 5 minutes.

### Competition assay

Overnight cultures of the SynCom members and the gfp-labelled *B. subtilis* (WT; *sfp* and *srfAC*) were pelleted (8000 rpm, 2 min) and resuspend in 0.1× TSB at an OD_600_ of 0.1. Next, 200 µL of a SynCom member was inoculated in the first row of 96-well microtiter plate. From there, the SynCom member was 10-fold diluted by transferring 20 µL of culture to the next row containing 180 µL of medium. This process was repeated for 6 dilution steps. Subsequently, 20 µL of the GFP-labelled *B. subtilis* variants was added to each well to establish the co-culture. Monocultures of both the SynCom member and *B. subtilis* variants served as controls to calculate competitiveness in co-culture. Cultivation was carried out in a Synergy XHT multi-mode reader (Biotek Instruments, Winooski, VT, US), at 25°C with linear continuous shaking (3 mm), monitoring the optical density and GFP fluorescence (Ex: 482/20; Em:528/20; Gain: 35) every 5 min. Kinetic parameters were estimated using the package GrowthCurver [93] in R. *B. subtilis* SMs induction by the SynCom spent media.

The WT strain was inoculated in the presence of culture spent media from the SynCom members. The spent media were obtained after 48h of growth in 0.1× TSB and filter through at 0.22 µm. 10% of the spent media to Erlenmeyer flasks containing potato dextrose broth (15 mL in 100 mL flasks), followed by inoculation with an overnight culture of P5_B1 (OD_600_ = 0.1). After 48 hours of incubation at 25°C and 220 rpm, the cultures were centrifuged, filtered, and subjected to HPLC analysis for surfactin detection.

### Assessment of *B. subtilis* establishment in diverse synthetic communities

To elucidate the role of surfactin in determining the establishment of *B. subtilis* within synthetic communities, we investigated whether P5_B1 can establish in various SynComs in a surfactin-dependent manner, using a methodology like the one described above for the competition assay. For this purpose, we selected five previously characterized bacterial SynComs, each with distinct compositions in terms of taxonomy and number of members, assembled for various objectives (Table S1). In all cases, the SynCom members and the gfp-labelled *B. subtilis* strains (WT and *srfAC*) were cultured overnight in 0.5x TSB. Following two washes with 0.9% NaCl, the ODs were adjusted to 0.1 in 0.1x TSB. The SynCom members were mixed in a 1:1 ratio and then inoculated and diluted in a 96-well plate. Subsequently, 20 µL of the gfp-labelled *B. subtilis* variants were added to each well to create the co-culture (Fig S10). Monocultures of both the SynCom member and *B. subtilis* variants were included as controls to determine competitiveness in the co-culture. Cultivation conditions and data analysis were conducted as described for the competition assay. Each experiment was performed with at least three independent replicates per treatment.

### Statistical analysis

Data analysis and graphical representation were performed using R 4.1.0 [94] and the package ggplot2 [95]. Statistical differences in experiments with two groups were explored via Student’s *t-*tests. For multiple comparisons (more than two treatments), one-way analysis of variance (ANOVA) and Tukey’s honestly significant difference (HSD) were performed. In all the cases, normality and equal variance were assessed using the Shapiro - Wilks and Levene test, respectively. Statistical significance (α) was set at 0.05. Detailed statistical analysis description for each experiment is provided in figure legends.

## Data availability

Analysis scripts, raw and processed data have been deposited at Github (https://github.com/carlosneftaly/SurfactinSynCom_story). Raw sequence reads of the RNAseq campaign have been deposited at the Sequencing Read Archive (SRA) with BioProject ID PRJNA1145146. LC-MS data have been deposited at GNPS-MassIVE under MSV000094405.

## Acknowledgment

The authors thank Morten D. Schostag for his suggestions on RNA analysis, Aaron J.C. Andersen and the DTU Bioengineering Metabolomics Core for support with LC-MS. This project was supported by the Danish National Research Foundation (DNRF137) for the Center for Microbial Secondary Metabolites. Funding from Novo Nordisk Foundation for the infrastructure “Imaging microbial language in biocontrol (IMLiB)” (NNFOC0055625) and the INTERACT project of the Collaborative Crop Resiliency Program (NNF19SA0059360) is acknowledged.

## Author contributions

Designed research: CNLA, ÁTK; performed the experiments: CNLA, CD, NAB; performed the chemical detection and analysis: MW, SJ; contributed analysis method: VHT, MLS; analyzed data: CNLA, MW, SJ; wrote the manuscript: CNLA, ÁTK with corrections by co-authors.

## Supplementary Information

**Fig S1.**
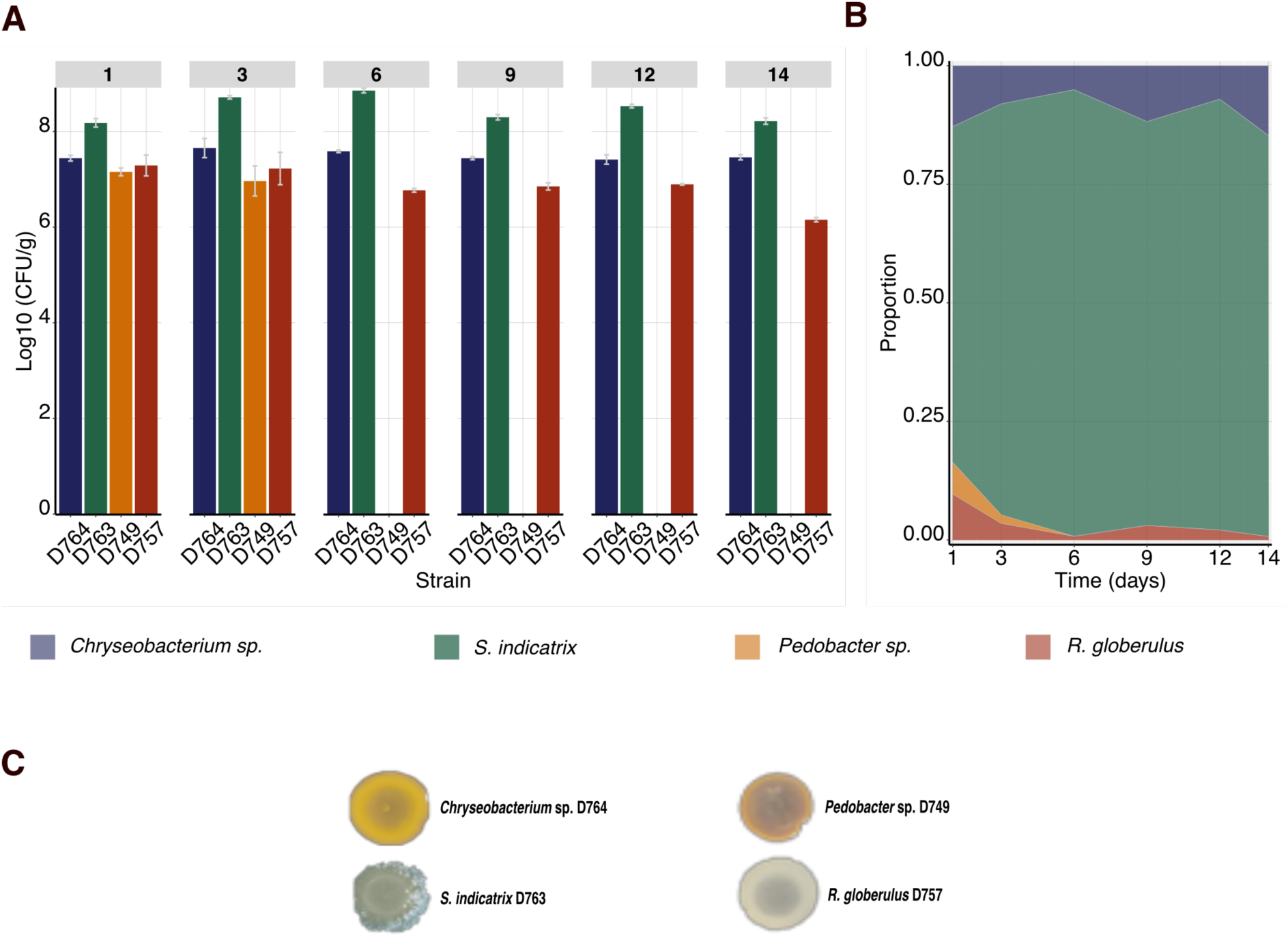
A stable, trackable simplified bacterial synthetic community (SynCom) for studying chemical ecology. Overnight cultures of the four members were mixed together and propagated in the hydrogel matrix. The abundance of each member was differentially determined as colony forming units per gram of beads based on their characteristic colony morphologies. (A). Population dynamics of each member over the time. (B) Members abundances represented as proportion occupied by each member related to the total biomass in the system. (C) Colony morphology of the SynCom members that allow us to differentially estimate their population in co-cultures. Error bars indicate standard deviation (n=3).

**Fig S2.**
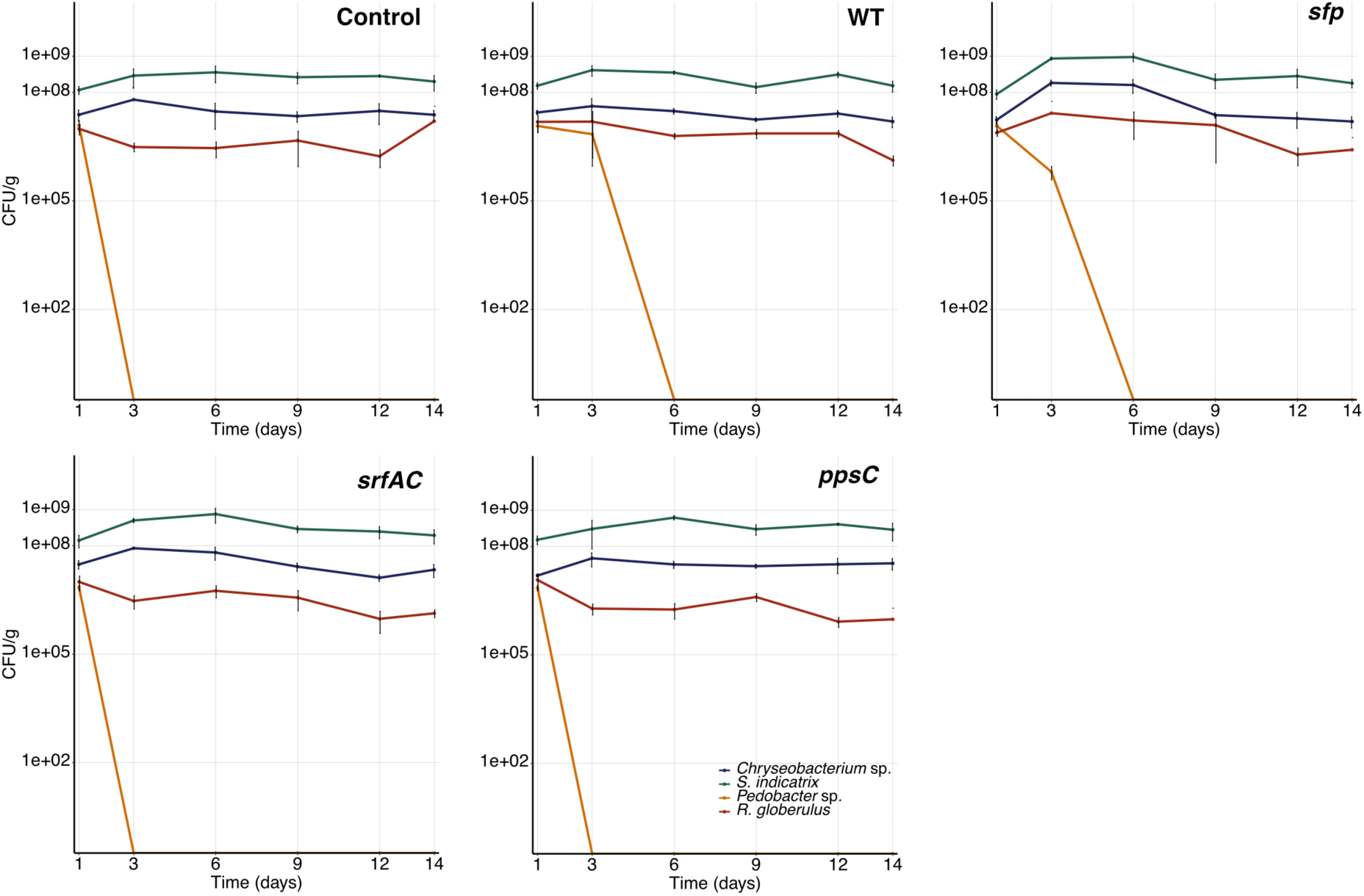
Raw counts (CFU/g) of each SynCom member when challenged with the different *B. subtilis* variants. These raw counts were used to generate the Fig1 A. Each line represents a SynCom member and the point the mean and the standard deviation (n=3).

**Fig S3.**
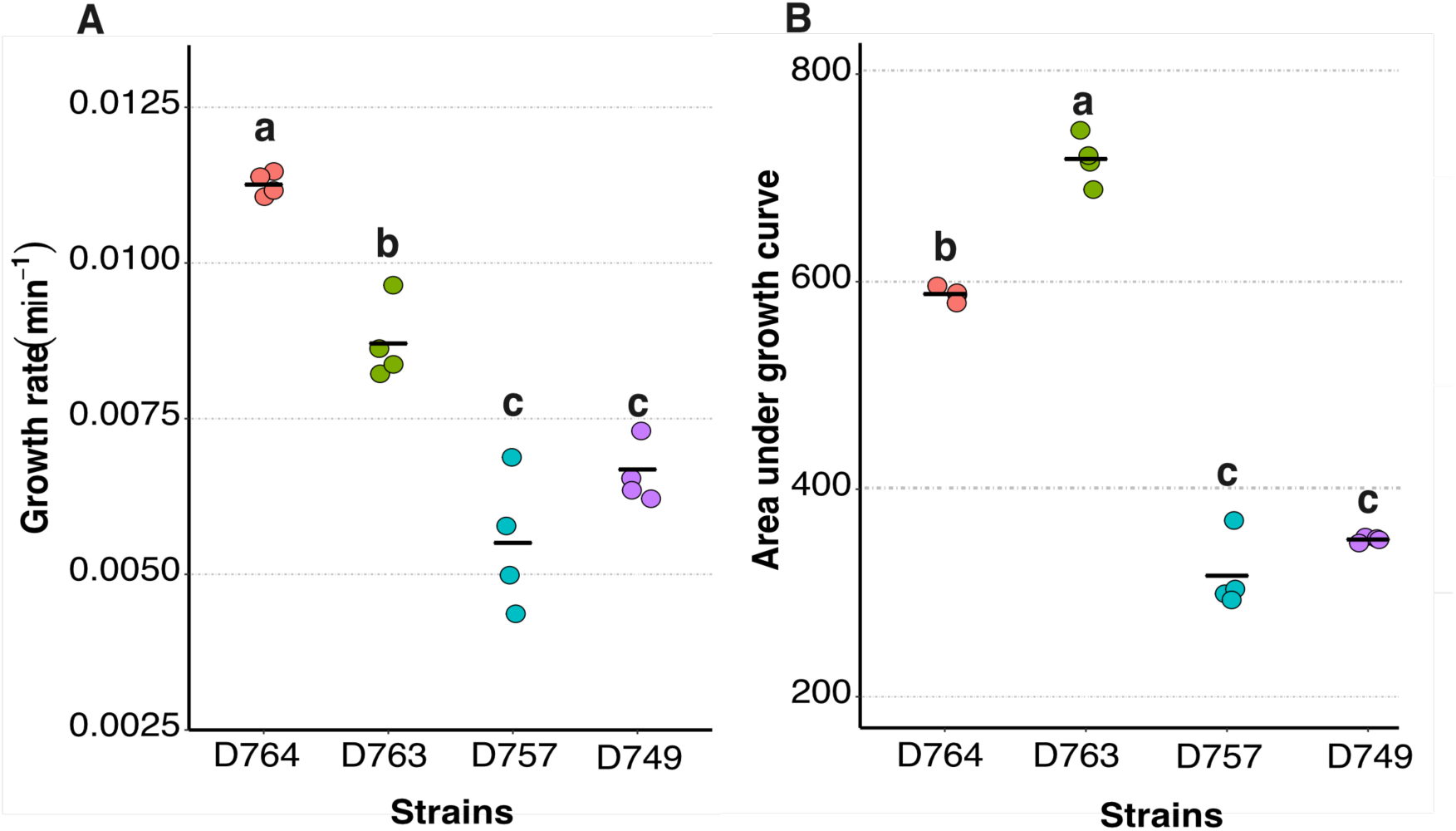
Growth properties of the SynCom members in liquid culture. The SynCom members were grown in 96-well microplates containing 0.1 × TSB at 25°C and 225 rpm for 24 h. Subsequently, the growth kinetic parameters were estimated using Growthcurver in R. *Chryseobacterium* sp. D764 and *S. inidcatrix* D763 are the fittest under these conditions as the growth rate and the area under the growth curve (AUDGC) revealed. (A) Growth rate of the SynCom members. (B) Area under growth curve as measure of strain productivity at the end of experiment. Data were derived from four replicates. Each point represents the samples and the black bar the mean of the group. Significant differences were evaluated using ANOVA and the letters indicate significant differences from a Tukey test (*P*<0.05).

**Fig S4.**
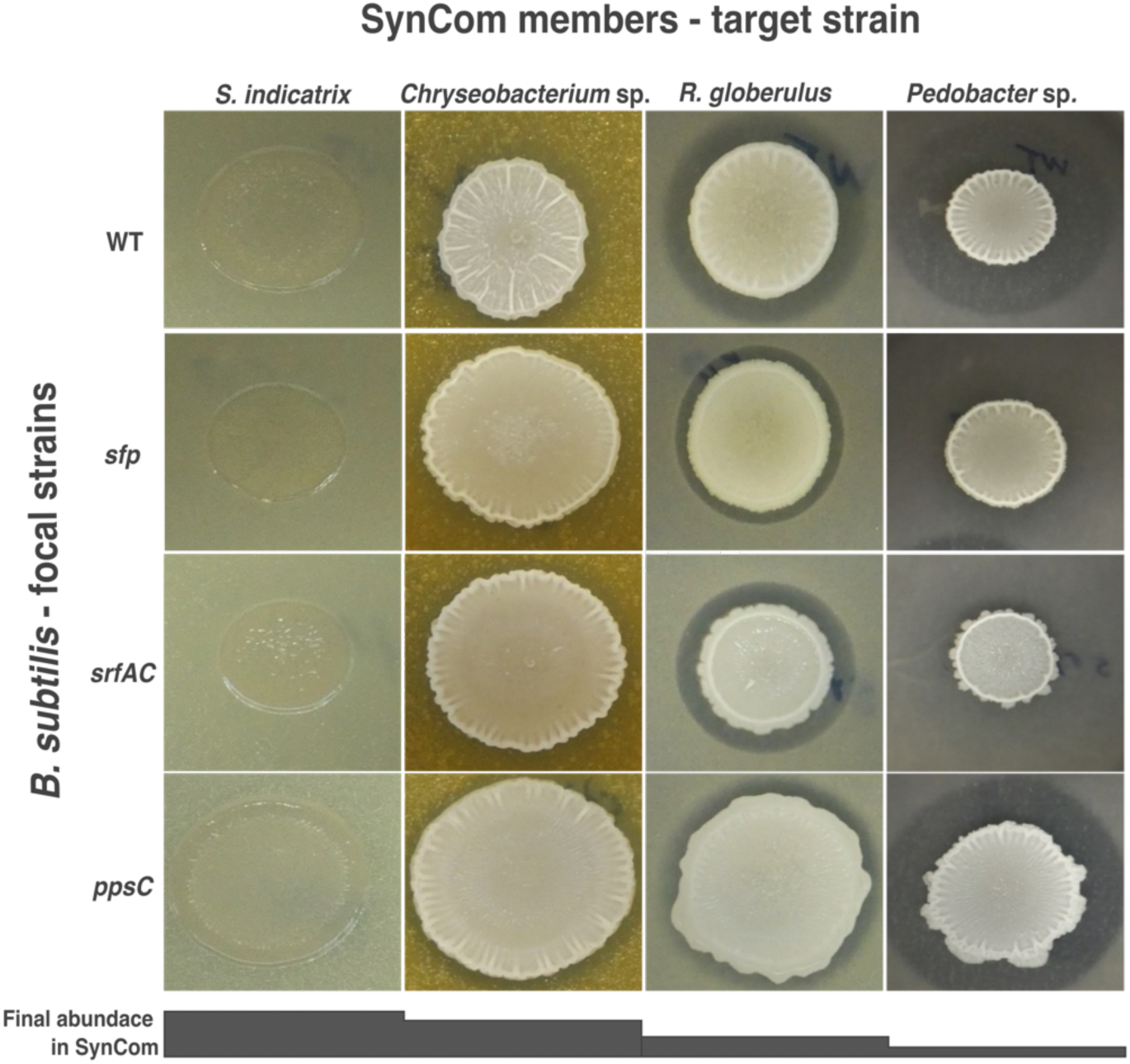
Testing the potential antagonistic activity of *B. subtilis* strains against SynCom members. The SynCom members were inoculated in soft layer of 0.1× TSA forming a lawn, on top 5µL of each *B. subtilis* strains was spotted. Then, the plates were incubated, and inhibition halos were recorded. The appearance of clearance halo indicates antagonistic activity. *R. globerulus* was susceptible to *B. subtilis* but such inhibition was not linked to NRPs production. In contrast, the inhibitory activity towards *Pedobacter* sp. was NRP-dependent, being surfactin crucial in such activity.

**Fig S5.**
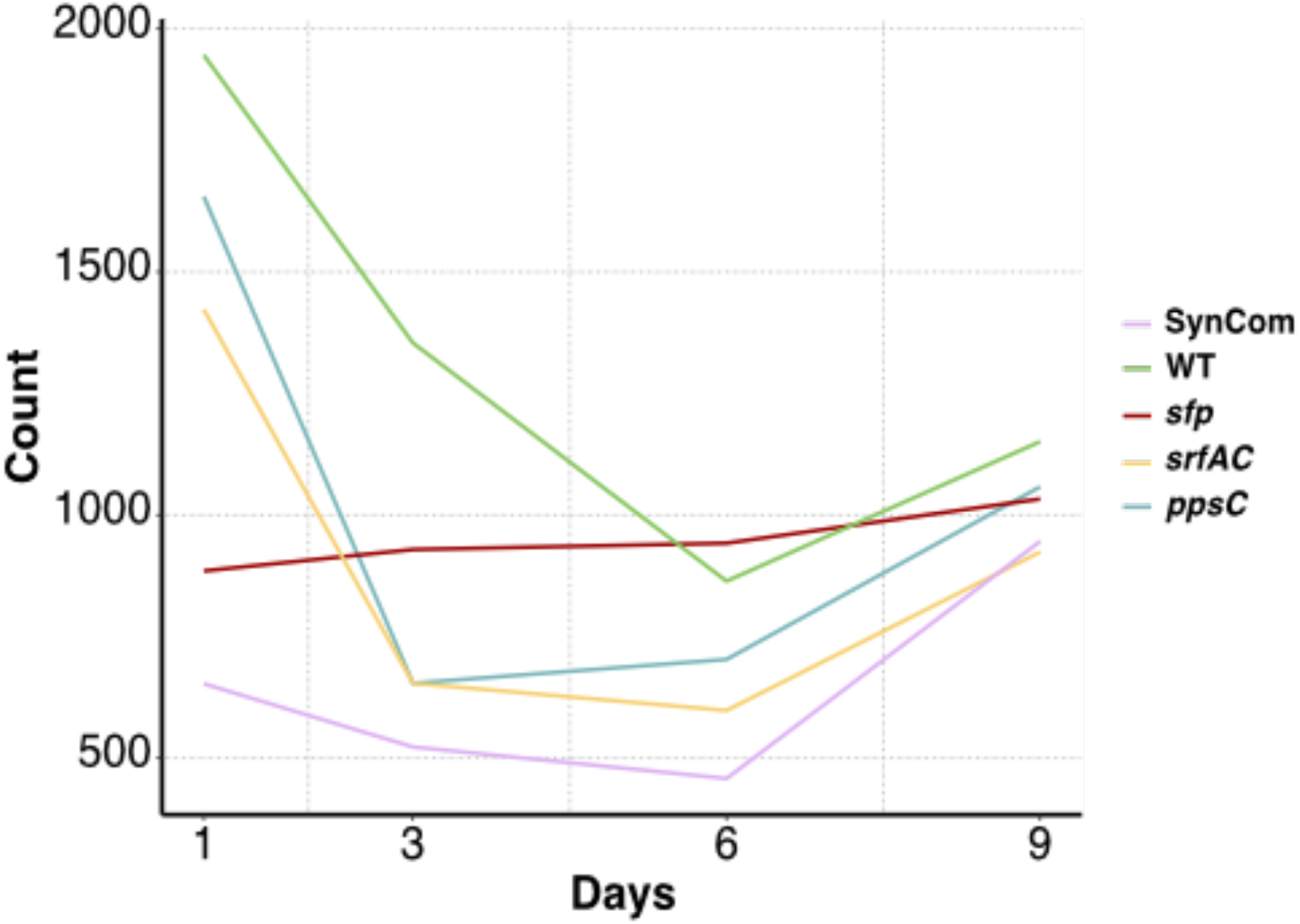
Temporal changes in the overall chemical output in the *B. subtilis* – SynCom co-cultivation experiment. The number of features detected in the different co-cultures were estimated from the ESI-MSI chromatogram and plotted over the time.

**Fig S6.**
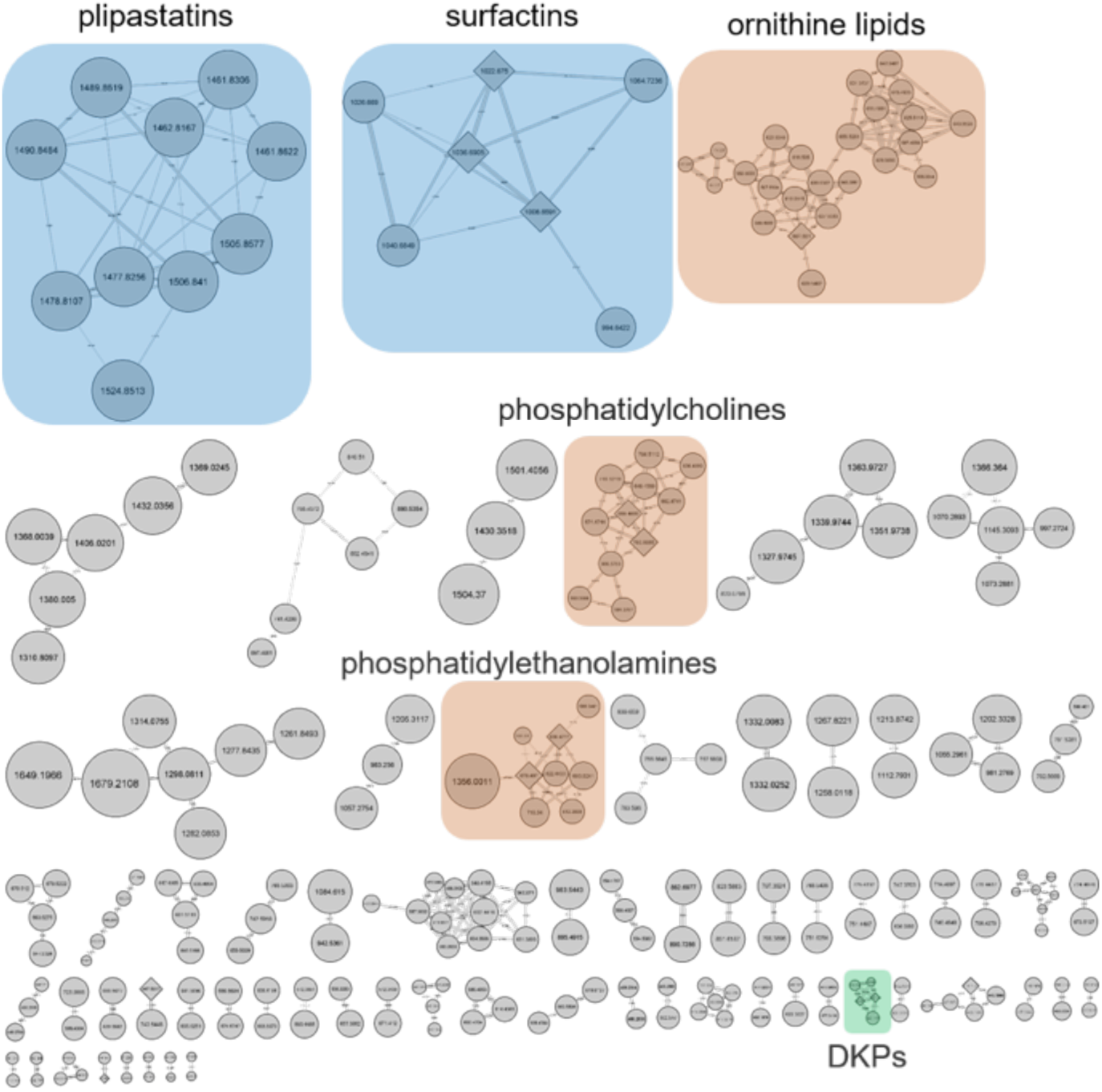
Feature-based molecular networking of the SynCom challenged with different variants of *B. subtilis.* Molecular networks of all detected features are presented in the plot. Identified chemical classes are highlighted on colors.

**Fig S7.**
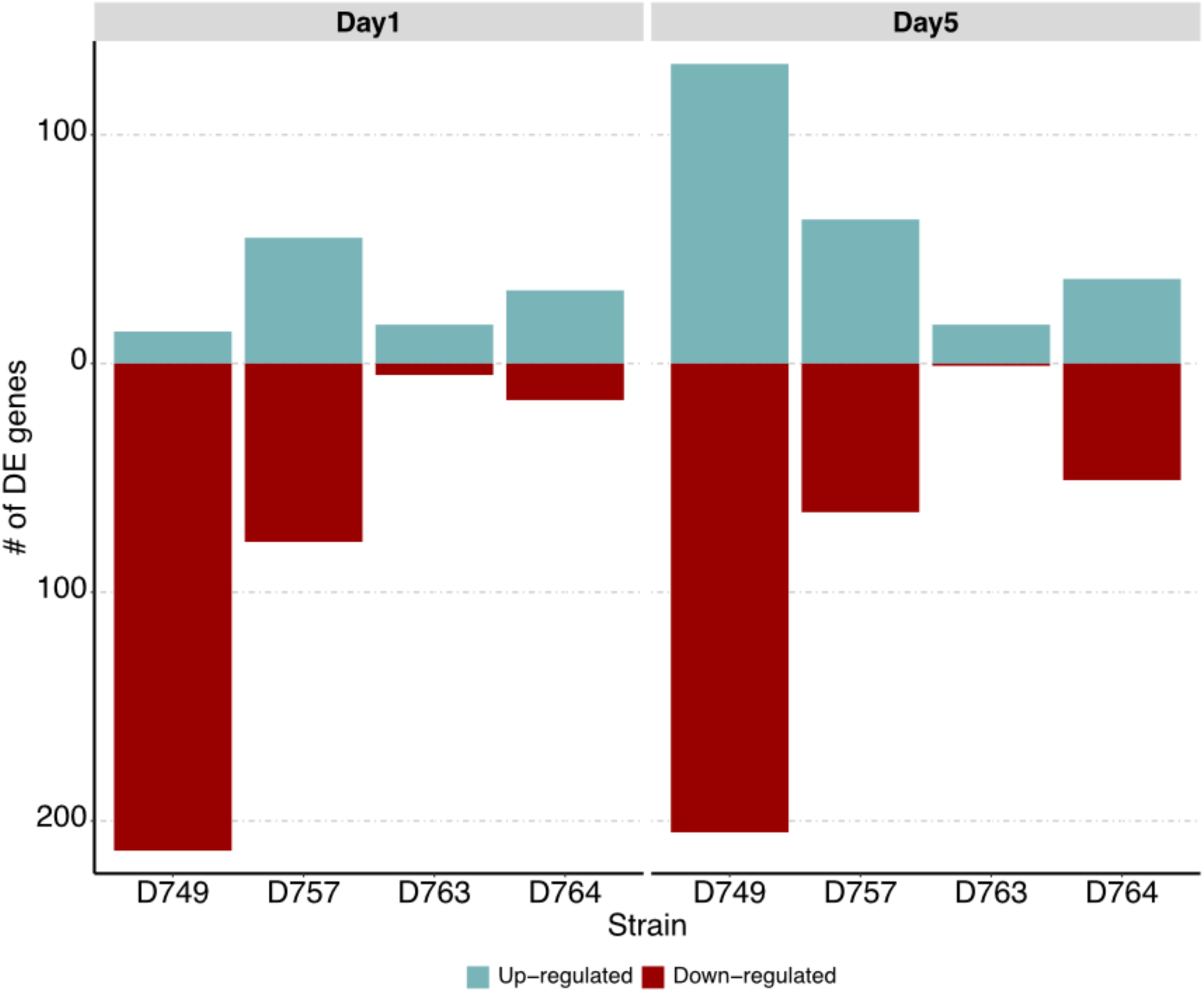
An overview of the differentially expressed genes of each strain over the time. Number of differentially regulated genes (log2FC ≥ |2| and p-value ≤ 0.01) in each species during co-cultivation experiment. In all the cases the comparisons were performed as WT vs. s*fp* mutant.

**Fig S8.**
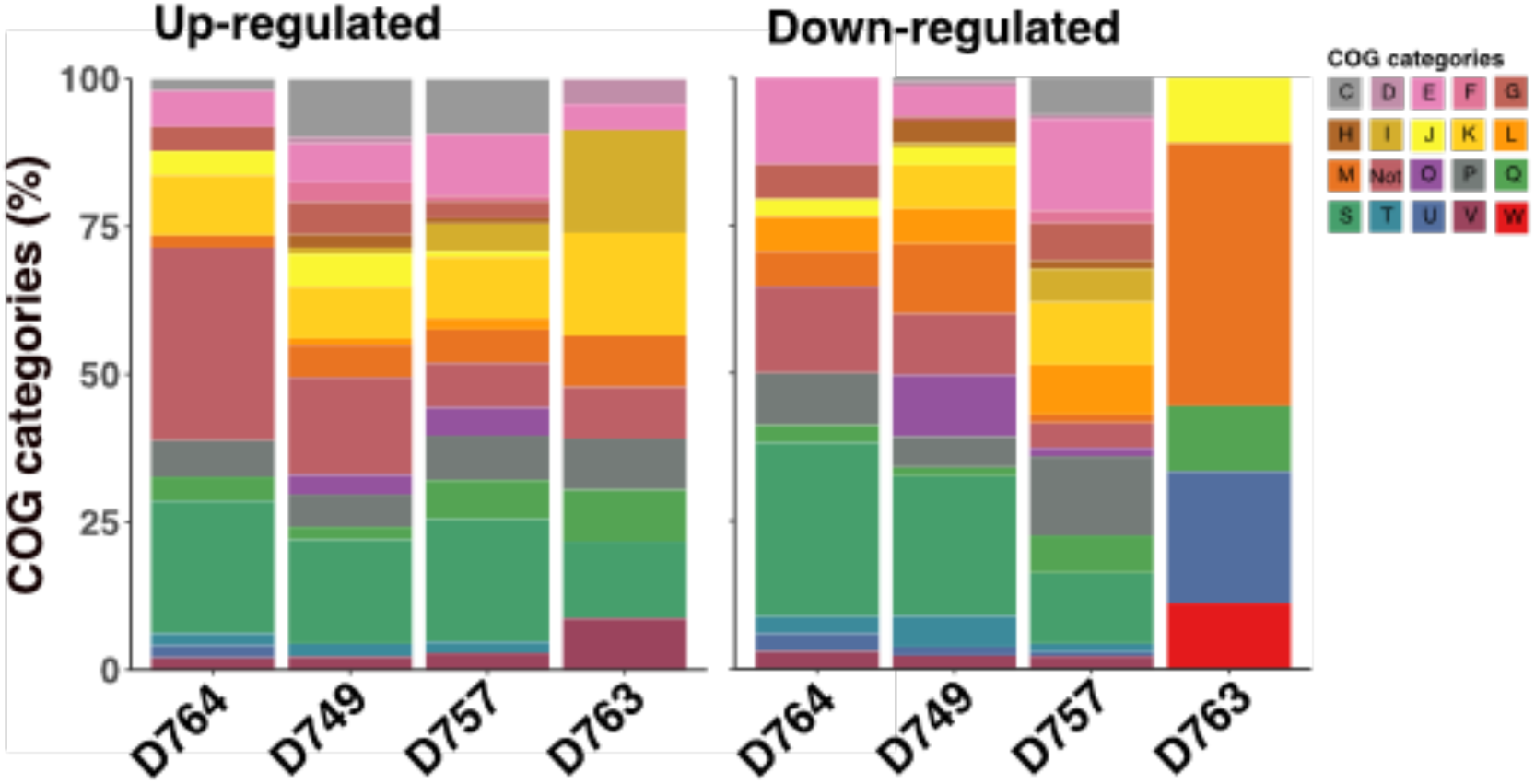
COG categories of genes up-or down-regulated by the four species in the SynCom challenged with the WT strain if compared to *sfp* ones at day 5 of sampling.

**Fig S9.**
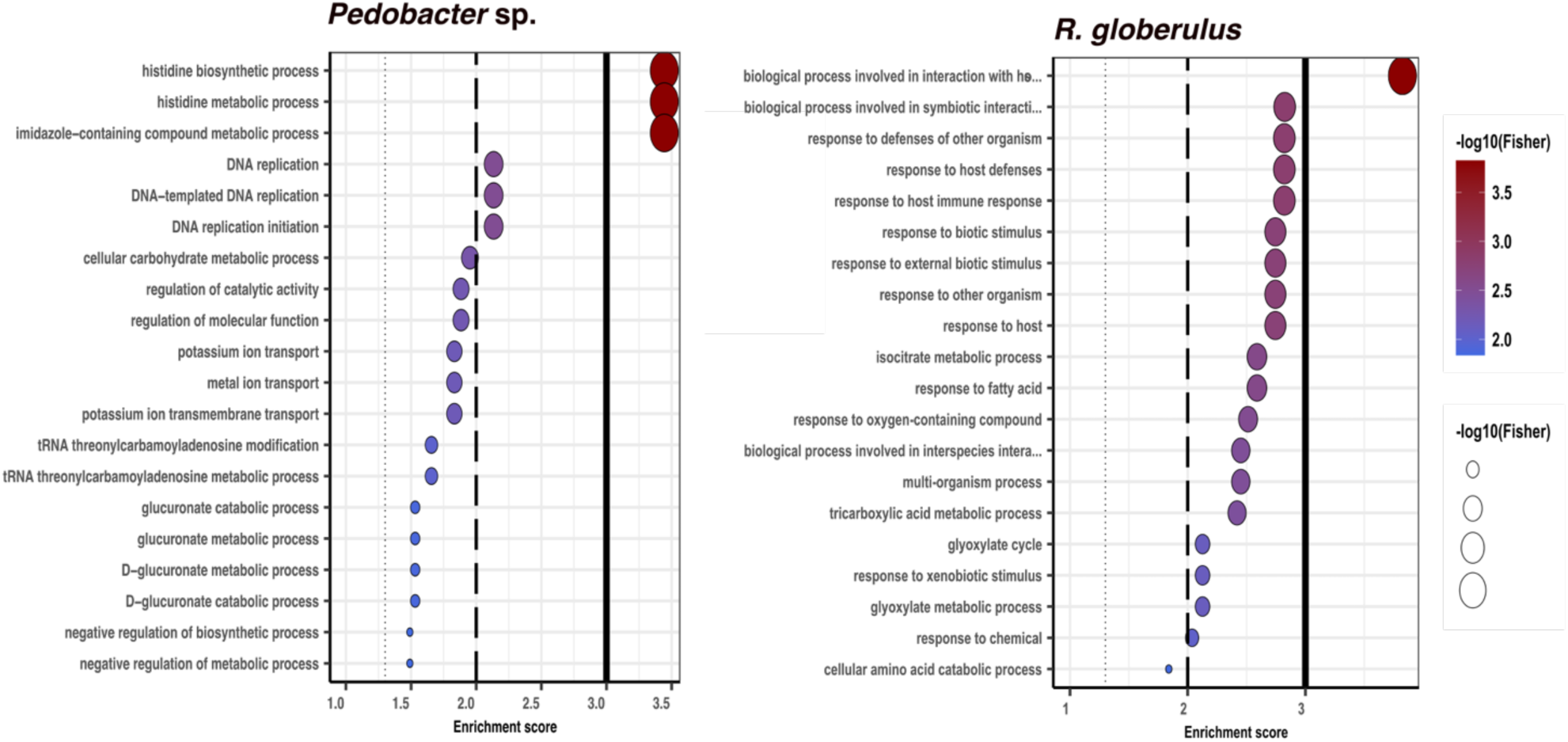
Biological processes in gene ontology (GO) enrichment analysis of DEGs during the *B. subtilis* – SynCom co-cultivation experiment. GO enrichment analysis was performed using topGO program in R. Only significantly enriched terms with corrected P < 0.05 were indicated. The color and size of each point represented the -log10 (FDR) values and enrichment scores. A higher -log10 (FDR) value and enrichment score indicated a greater degree of enrichment. **ynCom co-cultivation experiment. The number of features detected in the different co-cultures were estimated from the ESI-MSI chromatogram and plotted over the time.**

**Fig S10.**
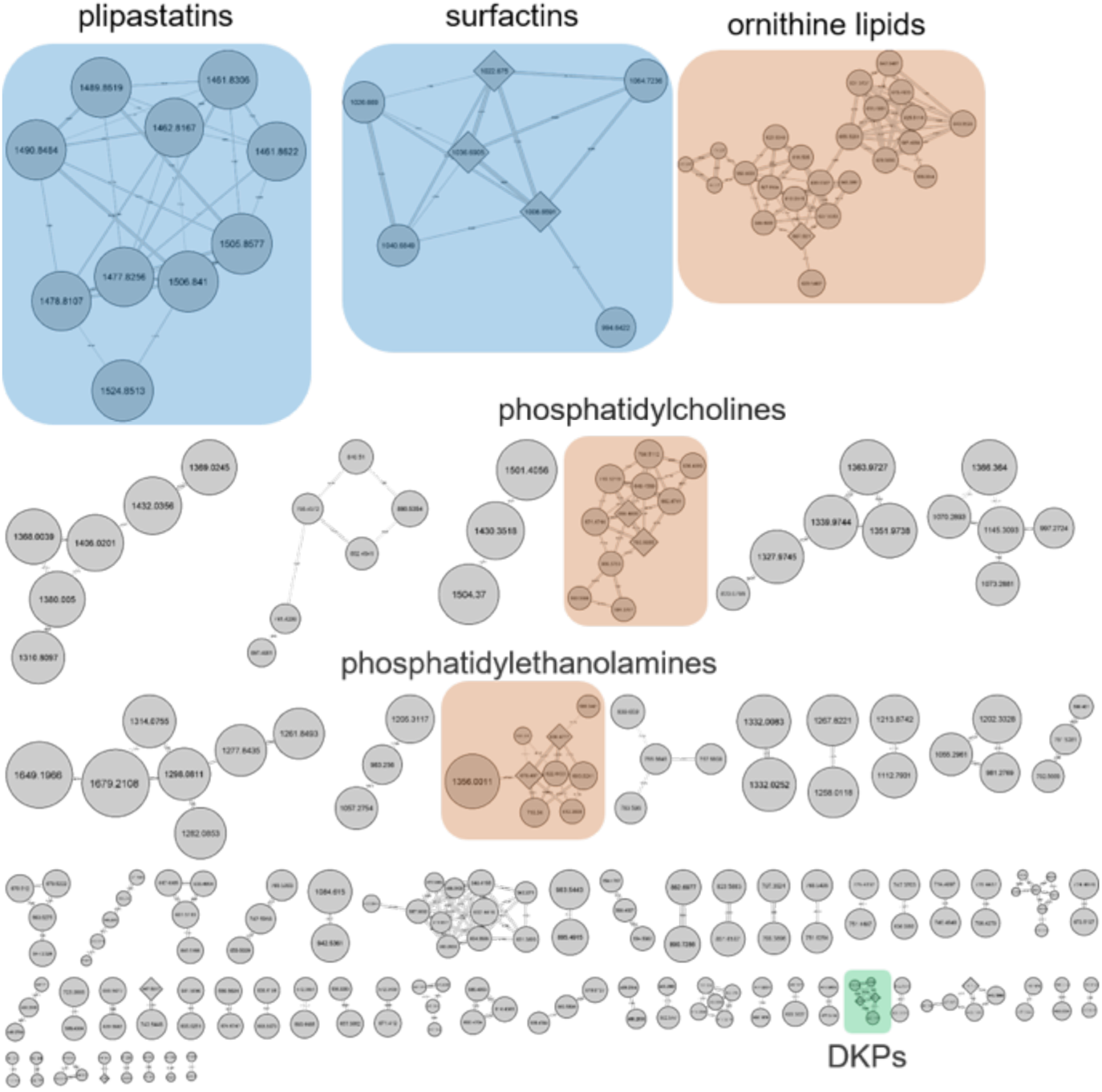
Feature-based molecular networking of the SynCom challenged with different variants of *B. subtilis.* Molecular networks of all detected features are presented in the plot. Identified chemical classes are highlighted on colors.

**Fig S11.**
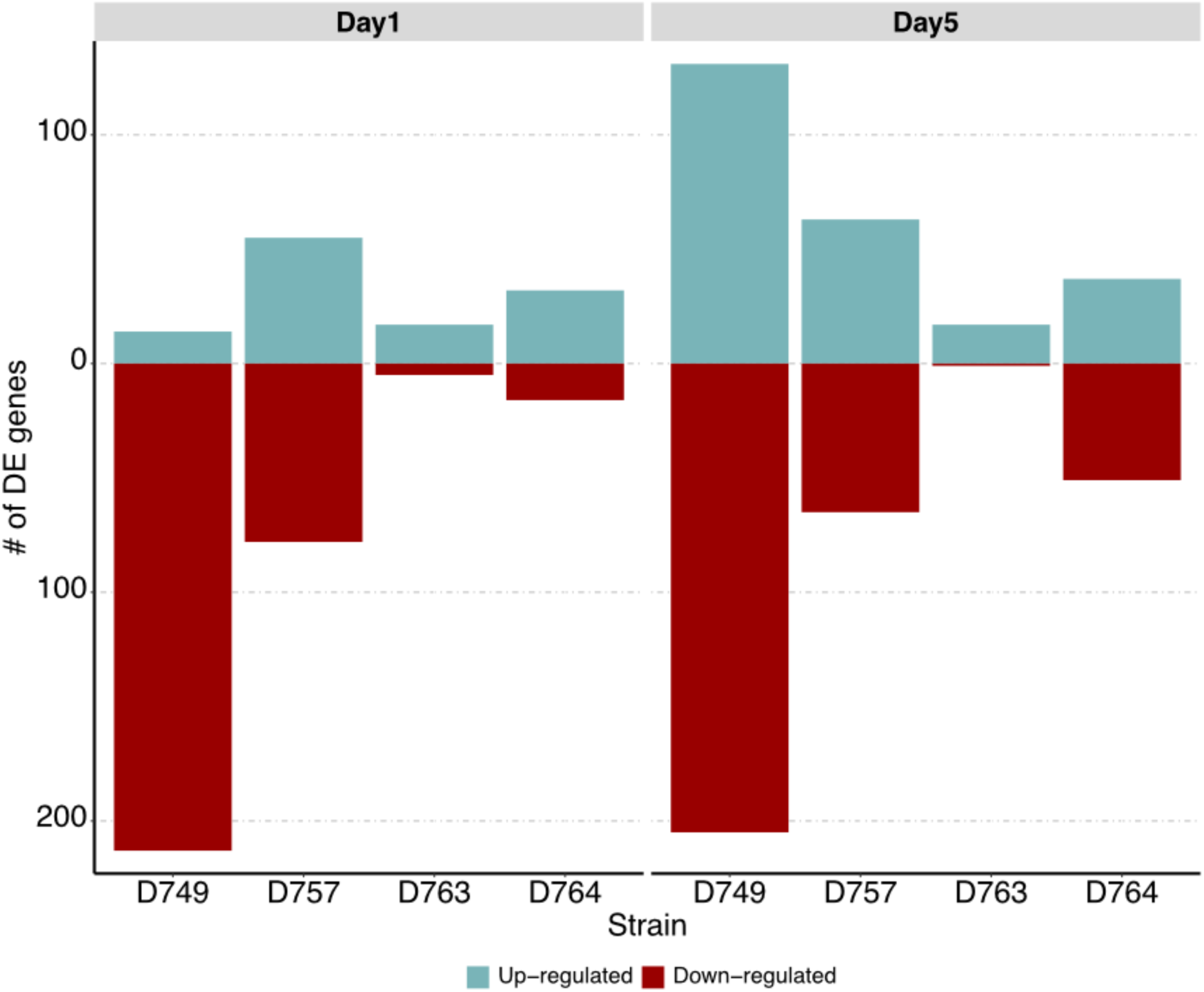
An overview of the differentially expressed genes of each strain over the time. Number of differentially regulated genes (log2FC ≥ |2| and p-value ≤ 0.01) in each species during co-cultivation experiment. In all the cases the comparisons were performed as WT vs. s*fp* mutant.

**Fig S12.**
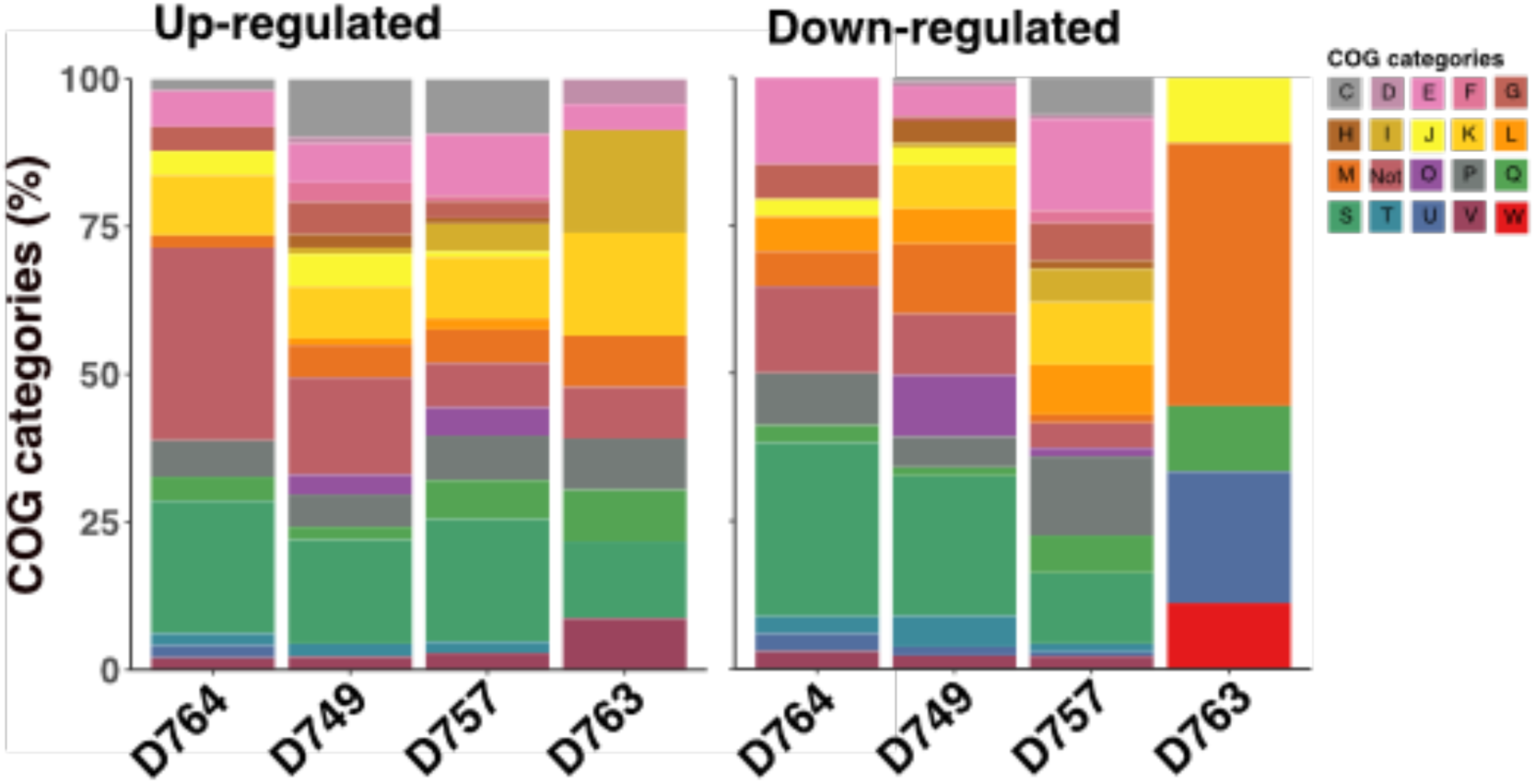
COG categories of genes up- or down-regulated by the four species in the SynCom challenged with the WT strain if compared to *sfp* ones at day 5 of sampling.

**Fig S13.**
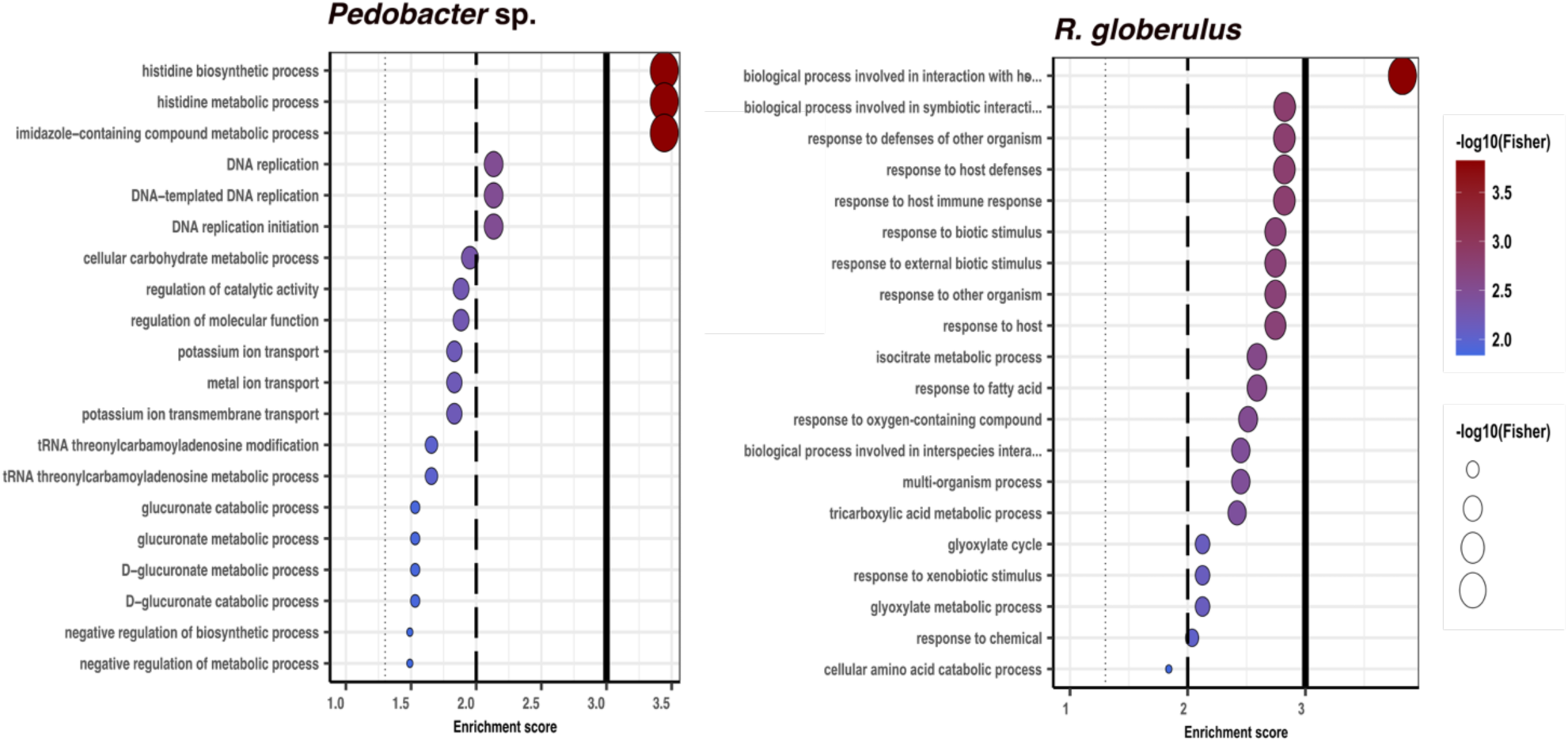
Biological processes in gene ontology (GO) enrichment analysis of DEGs during the *B. subtilis* – SynCom co-cultivation experiment. GO enrichment analysis was performed using topGO program in R. Only significantly enriched terms with corrected P < 0.05 were indicated. The color and size of each point represented the -log10 (FDR) values and enrichment scores. A higher -log10 (FDR) value and enrichment score indicated a greater degree of enrichment.

